# High-stability, clamp-free soluble Sarbecovirus spike trimers and their potential for pan-Sarbecovirus vaccine development

**DOI:** 10.1101/2025.09.14.676173

**Authors:** Christine Langer, Irene Boo, Tasnim Zakir, Charley Mackenzie-Kludas, Johnson Samuel, Stewart A. Fabb, Chee Leng Lee, Rob J. Center, Anupriya Aggarwal, Stuart Turville, Colin Pouton, Kanta Subbarao, Fasséli Coulibaly, Heidi E. Drummer, Pantelis Poumbourios

## Abstract

Broadly effective vaccines are needed to protect against future pandemics caused by severe acute respiratory syndrome coronavirus (SARS CoV)-like coronaviruses (sarbecoviruses). The development of simple trimeric subunit vaccines based on the Sarbecovirus spike (S) has proven problematic due to the unstable nature of the S trimer. Here we developed clamp-free, highly stable soluble S trimers by truncating the stem helix to maximize yield and covalently linking the 3 monomers via engineered disulfides to increase thermostability. In K18hACE2 mice, covalently linked SARS CoV-2 S trimers elicited >10-fold higher neutralizing antibody (NAb) titres than parental unlinked trimers and protected the mice against viral challenge. A trivalent vaccine formulation comprised of covalently stabilized spikes derived from 3 divergent ACE2-using Sarbecovirus clades elicited broad and potent neutralizing activity in mice. The covalently linked S trimers were stable at 37°C for 112 days and remained intact following lyophilization and storage at ambient temperature for 6 months. This study establishes a framework for producing simple and stable highly immunogenic pan-Sarbecovirus S subunit vaccines that can be stored and distributed in the absence of a cold chain.

---

Coronaviruses (CoVs) infect more than 500 bat and other animal species, which serve as reservoirs for CoV evolution and potential sources of spillover into humans ^1-3^. Two such spillover pathogens, SARS CoV and SARS CoV-2 caused ∼8,000 and more than 700,000,000 reported human cases, respectively, with significant morbidity and mortality. SARS CoV and SARS CoV-2 are Sarbecoviruses, a subgenus of the βCoVs, and are classified into 4 clades; clade 1a (e.g. SARS CoV), clade 1b (e.g. SARS CoV-2), and clade 3 viruses use ACE2 as the entry receptor, whereas clade 2 viruses use an alternative receptor ^4-6^. The transmission of a novel animal Sarbecovirus to humans could result in another pandemic. Vaccines that provide broad protection against this diverse reservoir of human pathogens are urgently needed.

The trimeric CoV spike (S) glycoprotein is responsible for receptor attachment and virus cell-membrane fusion that leads to viral entry. The spike is comprised of 2 non-covalently linked subunits: S1, which contains the receptor binding domain (RBD) and S2 which contains the fusion peptide and membrane anchor (for review, see: ^7,8^. S is the target of protective NAbs and for SARS CoV-2 is the basis of successful vaccines. For example, full-length S delivered either as mRNA, protein nanoparticle or viral vectored vaccines has been estimated to have saved over 14 million lives ^9^. These vaccine modalities can be applied in principle to new pandemics; however, the cold chain requirements of mRNA vaccines, the complexity and high cost of protein nanoparticle vaccines and the adverse events associated with viral vectored vaccines has limited access and contributed to global inequities. A simple method for producing conventional subunit vaccines that can be stored and distributed independently of a cold chain and can generate broad immunity against Sarbecoviruses is needed for the prevention of future pandemics. To this end, a soluble and stable form of the S trimer has been developed by replacing the membrane spanning sequence with a trimerization clamp derived from foreign proteins. However, such clamps can themselves elicit off-target antibody responses that can hamper passage of new vaccines through the developmental/regulatory pathway ^8,10-13^.

The aim of this study was to produce clamp-free, intrinsically stable soluble S trimers with improved biochemical antigenic and immunogeic characteristics. A trivalent vaccine formulation comprised of clamp-free stabilized spikes derived from 3 divergent Sarbecovirus clades elicited broad and potent neutralizing activity in mice. This study establishes a framework for producing simple and stable highly immunogenic pan-Sarbecovirus S subunit vaccines that can be stored and distributed in the absence of a cold chain for pandemic prevention.

## Results

### The stem region controls S oligomer yield and thermostability

The A1016V/A1020I ‘VI’ mutation in the central helix of S2, which fills a cavity in the SARS CoV-2 S trimer core, enabled the production of stable clamp-free trimers ^14^. The stem/HR2 region of this construct (here called S2P.BA5.VI-1208) (**Figure. 1a)**, was sequentially truncated **(Fig. 1b)** to identify determinants of trimer yield and thermostability. Size exclusion chromatography (SEC) of affinity-purified S2P.BA5.VI-1208 revealed heterogeneous protein; truncation to G1204, Q1201, and L1200 resulted in a more prominent putative trimer peak **(Fig. 1c)**. The melting temperatures (Tm) of purified trimers (**Fig. 1d)** were determined in the range 60°C-61°C by differential scanning fluorimetry (DSF) (**Fig. 1e**).

**Fig. 1.**
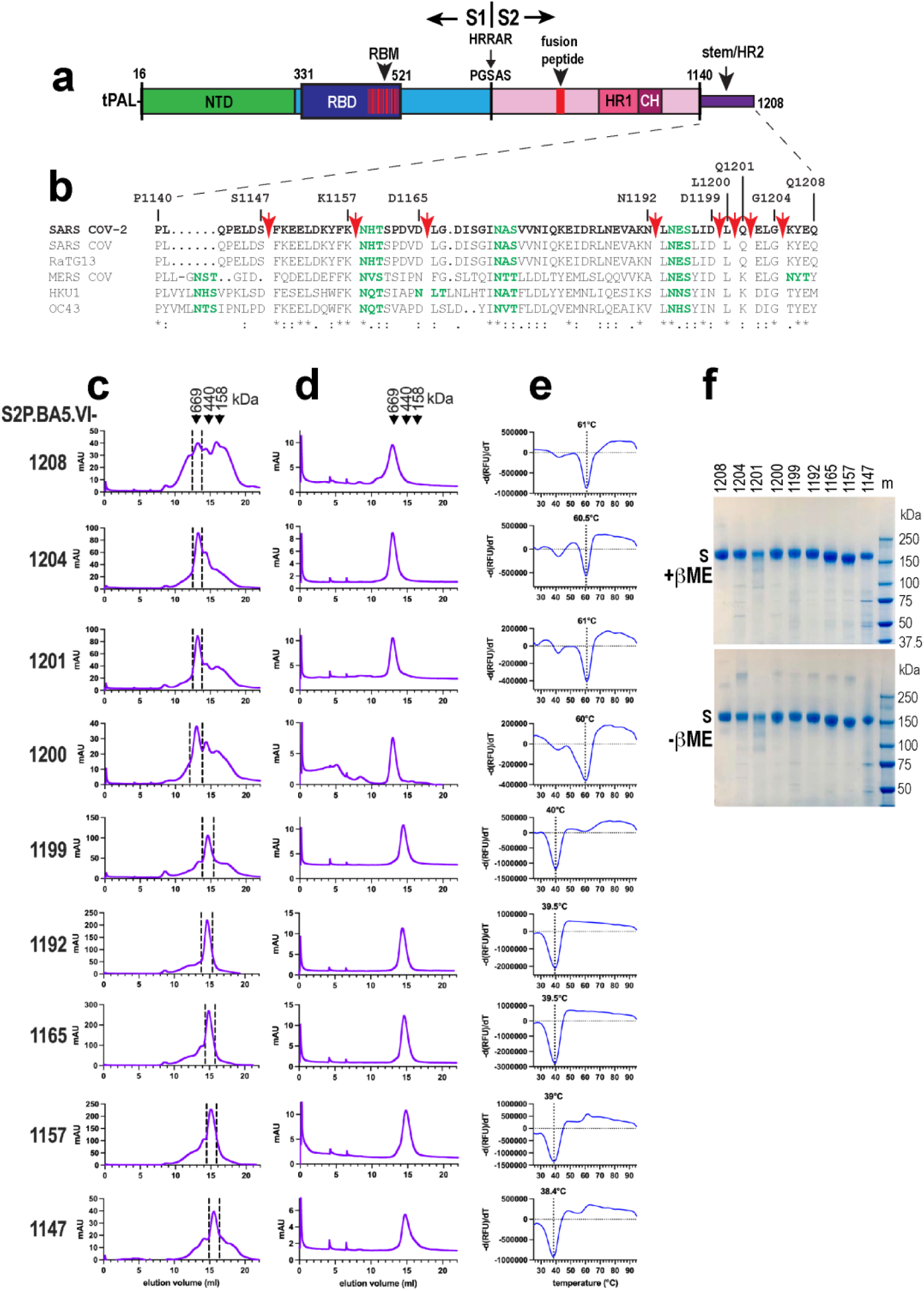
Biochemical characteristics of omicron BA.5 glycoproteins with truncated stems. **a.** Linear map of S2P.BA5-1208. tPAL: tissue plasminogen activator leader sequence, NTD, N-terminal domain; RBD, receptor binding domain; RBM, receptor-binding motif; HRRAR->PGSAS, furin site mutation, HR1, heptad repeat 1; CH, central helix, HR2, heptad repeat 2. **b.** Alignment of Betacoronavirus stem sequences. Red arrows indicate truncation points. * = identical amino acids; : and. = similar amino acids. Potential N-linked glycosylation sites in green. **c.** Superose 6 SEC of secreted spike proteins eluted from TALON resin. **d.** Fractions within the vertical dashed lines in **c** were pooled, concentrated and re-chromatographed on Superose 6 following a freeze (-80°C)-thaw cycle. **e.** Melting temperature of purified spike oligomers shown in **d** determined by DSF. **f.** SDS-PAGE of purified spike proteins shown in **d**. βME, betamercaptoethanol.

Further truncation to D1199, N1192, D1165, K1157 and S1147 gave rise to major peaks coeluting in SEC with the 440 kDa marker **(Fig. 1b,c,d)**. Their elution position indicated a smaller hydrodynamic radius ^15^ relative to the longer spikes, likely due to the removal of C-terminal stem residues that may form a mobile extended structure at the base of the trimer ^16^. These shorter S2P.BA5.VI proteins had lower Tms in the range 38.4°C-40°C **(Fig. 1e)**. SDS-PAGE indicated a prominent band at ∼160-170 kDa for the spike proteins (**Fig. 1f**). Plots of Tm and trimer yield as functions of C-terminal length show that L1200 is the key determinant of thermostability. However, the longer thermostable trimers are low yielding. Conversely, truncation to N1192, D1165 and K1157 resulted in high yielding S oligomers but with lower thermostability (**Supplementary** Fig. 1a).

### NAb epitope profiles of S2P.BA5.VI trimers with truncated stems

The NAb epitope profiles of the purified S2P.BA5.VI proteins were determined in biolayer interferometry (BLI) **(Supplementary** Fig 2). Plots of the responses at binding equilibrium (R_eq_) **(Supplementary** Fig 1b) shows roughly equivalent binding by all mutants to the NTD monoclonal NAb (mNAb) C1520. K1157-, D1165- and N1192-terminated proteins achieved higher R_eq_ values with receptor-binding motif (RBM)-directed ligands ACE2-Fc, Omi-18 and Omi-42 ^17^ than longer constructs. A similar trend was observed with the RBD flank-specific mNAbs, S2H97 and SP1-77 ^6,18^. Poor overall binding was observed with the fusion peptide mNAb COV44-79^19^, indicating that this epitope is poorly accessed.

S2P.BA5.VI-1192 exhibited maximal binding to the stem mNAb CV3-25 which has broad Sarbecovirus neutralizing properties ^20^. The binding kinetics (**Supplementary Table 1**) indicate relatively high affinity interactions (KDs≤6.7x10^-9^M) between all spike proteins and ligands (except COV44-79).

### Structure-directed cysteine substitution mutagenesis to covalently stabilize the S2P.BA5.VI trimer

In parallel with the above, structure directed Cys-substitution mutagenesis was used to stabilize the S2P.BA5.VI ectodomain trimer via engineered intermolecular disulfides. S1-S1, S1-S2 and S2-S2 interfacial residues were identified using the LPC-CSU server (http://oca.weizmann.ac.il/oca-bin/lpccsu) ^21^ for Cys replacement. Eleven contact residue pairs were subjected to Cys substitution mutagenesis in the context of S2P.BA5.VI-1147 (**Supplementary** Fig. 3a,b). Ser1147 was initially chosen as the terminating residue as it is the last amino acid observed in many cryo-EM structures of spike trimers (e.g. ^22,23^). D571C/N967C (D17), A570C/N967C (I1), and N914C/S1123C (L23) mutations increased the yield and Tm of S2P.BA5.VI-1147 (**Supplementary** Fig. 3c). D17 and I1 possessed the highest Tms: 55.5°C and 51.5°C, respectively (**Supplementary** Fig. 3e). Nonreducing SDS-PAGE revealed molecular weights >250 kDa for D17, I1 and L23, which were resolved to the expected monomer molecular weight (∼170-180 kDa) under reducing conditions, consistent with highly efficient disulfide formation (**Supplementary** Fig. 3f). The L23 mutant migrated faster than D17 and I1 mutants in nonreducing SDS-PAGE suggesting that 2 monomers/trimer were likely linked for L23 whereas 3 appear to be linked for D17 and I1. In SEC, the D17, I1 and L23 mutants exhibited a larger hydrodynamic radius than the S2P.BA5.VI-1147 parent (**Supplementary** Fig 3d).

### I1 and D17 mutations in N1192-truncated trimers

**Supplementary** Fig. 1b showed that S2P.BA5.VI-1192 exhibited superior neutralizing ligand reactivity. D17 and I1 mutations were therefore introduced to omicron BA.5 N1192-truncated constructs: S2P.BA5.VI-1192, S2P.BA5-1192 (lacking VI) and S6P.BA5-1192 constructs. The latter contains the ‘6P’ mutation (F817P, A892P, A899P, A942P, V986P, K987P) shown to improve yield and thermostability of foldon-clamped ancestral SARS CoV-2 spike ^24^. The TALON-purified spikes largely resolved as single peaks in SEC (**Fig. 2a)**. The highest yields were obtained with the ‘6P’ mutation (**Fig. 2e)**. The D17 and I1 purified trimers (**Fig. 2b)** had Tms of 55°C and 51°C, respectively, in the 3 spike contexts (**Fig. 2c**). Nonreducing SDS-PAGE indicated that virtually all of the D17 and I1 spike monomers within oligomers were disulfide linked (**Fig. 2d**). Secreted S2P.BA5.VI-1192.L23 protein was not detected therefore this mutant was not pursued (**Fig. 2e**).

**Fig. 2.**
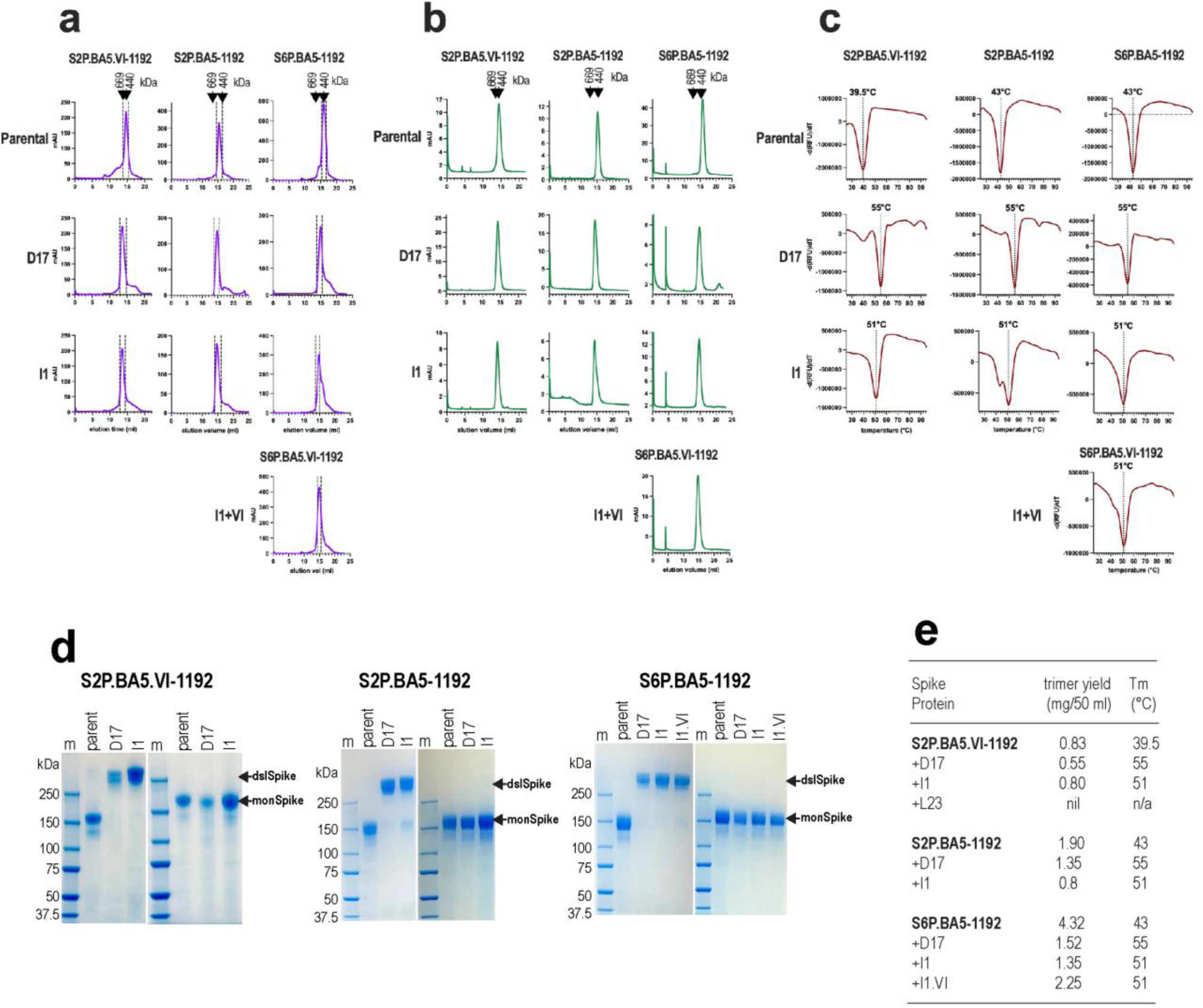
Biochemical characteristics of soluble covalently stabilized clamp-free omicron BA.5 S oligomers. **a.** Superose 6 SEC of secreted spike variants eluted from TALON resin. **b.** SEC of purified spike glycoproteins. The fractions within the vertical dashed lines in **a** were pooled, concentrated and then re-chromatographed following a freeze (-80°C)-thaw cycle. **c.** Melting temperature of purified S oligomers determined by DSF. **d.** SDS-PAGE of purified S oligomers. dslSpike, disulfide-linked spike; monSpike; spike monomer. **e.** Yield and Tm of purified omicron BA.5 spike oligomers.

To determine whether D17 and I1+VI overcome the need for 2P and 6P ^23,24^ for S trimer expression and stability, Pro-substituted residues were reverted to the native amino acids (SnoP constructs). SnoP.D17-1192 and SnoP.I1+VI-1192 trimers represented ∼50-70% of total secreted spike (**Supplementary** Fig. 4a). The purified oligomers (**Supplementary** Fig. 4b) had identical Tms to their 2P-and 6P-containing counterparts (**Supplementary** Fig. 4c) but were obtained in lower yields. SDS-PAGE under non-reducing conditions of SnoP oligomers indicated quantitative disulfide bonding between monomers within oligomers (**Supplementary** Fig. 4d). The 2P and 6P mutations are therefore not essential for trimer stability nor D17 and I1 disulfide formation but do promote high trimer yields of the VI containing construct.

BLI was used to assess the presentation of mNAb epitopes in the S6P.BA5-1192 series (**Supplementary** Fig. 5**; Supplementary Table 2**). Parental S6P.BA5-1192 showed robust binding to RBM region ligands (ACE2-Fc, Omi-18, Omi-42 and SA55), whereas D17 and I1+VI versions exhibited diminished binding.

S2H97, SP1-77 and S309, directed to the conserved flanks of the RBD, bound with similar potency to the 3 constructs. The data suggest that the RBM region is somewhat occluded in D17 and I1+VI oligomers while conserved regions of the RBD remain exposed. This idea is illustrated by **Supplementary** Fig. 6 showing that the epitopes of RBM-directed mNAbs are exposed in a trimer with an RBD in the ‘up’ orientation but occluded when all 3 RBDs are ‘down.’ By contrast, conserved epitopes on the RBD flanks are exposed in both conformations.

Cryo-EM of the omicron BA.2.86 homolog of S6P.BA5-1192.D17 (S6P.BA286-1192.D17) provided further evidence for the RBD-down ‘closed’ conformation. (The biophysical and antigenic properties of S6P.BA286-1192.D17 are very similar to its BA.5 counterpart [see Fig. 4 and Supplementary Fig. 11]). A typical pre-fusion spike trimer with an arrangement of RBD heads in the down orientation is observed in **Supplementary** Fig. 7e. We note that due to a preferred spike orientation on EM grids, the reconstruction does not allow modelling but a fit of the PDB ID 6X2C spike structure ^25^ suggests that the RBM and RBM-overlap epitopes will be occluded, while epitopes on the flanks of the RBD remain exposed as indicated in BLI experiments.

The pan-variant-neutralizing stem mNAb, CV3-25, exhibited markedly diminished off-rates with D17 and I1+VI mutants relative to parental S6P.BA5-1192, indicating that the D17 and I1 mutations stabilize stem-CV3-25 interactions (**Supplementary** Fig. 5**; Supplementary Table 2**).

### The core-filling VI mutation cooperates with L1200 to confer thermostability to clampless spike trimers

We previously showed that VI was necessary to maintain the quaternary structure of the clamp-free S2P.BA5-1208 spike ^14^ but the data presented in Fig. 2 indicated that is not the case for N1192-terminated constructs. We therefore asked whether VI operates in conjunction with the stem amino acid L1200 that confers higher thermostability to clamp-free trimers (**See Fig. 1; Supplementary** Fig. 1a). VI was reverted to A1016/A1020 in S2P.BA5-1200 and SEC-purified oligomers were subjected to 2 freeze-thaw cycles, each followed by SEC and DSF. Whereas S2P.BA5.VI-1200 eluted as a single ∼669 kDa species with a Tm of 60.5°C following all treatments, increasing amounts of a lower mol.wt species appeared after the first and second freeze-thaw cycles for its AA-reverted counterpart (**Supplementary** Fig. 8a,b). Furthermore, each AA-reverted S2P.BA5-1200 freeze-thaw cycle was associated with increasing amounts of a thermolabile 42°C-Tm species. These data suggest an allosteric connection between the VI mutation in the core of the spike trimer and L1200 in the stem which confers oligomer stability to constructs with longer stem sequences.

### Immunogenicity and protective effect of S6P.BA5-1192 oligomers in K18hACE2 mice

Four groups of K18hACE2 mice (n=16) received 3 doses of vehicle (PBS) or 10 μg parental, D17- or I1+VI-mutated S6P.BA5-1192 oligomers in AddaVax adjuvant. (S6P.BA5-1192.I1+VI was included as it is the highest yielding covalently linked trimer). The mice were bled immediately before the 3^rd^ dose and 2 weeks after the 3^rd^ dose (**Fig. 3a**). The R-20 microneutralization assay ^26^ revealed that mean homologous virus neutralizing ID50s were modestly boosted (3.3-4.6-fold) by the 3^rd^ immunization (**Supplementary** Fig. 9a). Analysis of post-dose 3 sera indicated that >10-fold higher homologous omicron BA.5 neutralizing ID50s were elicited by the D17 and I1+VI immunogens (geometric mean ID50 = 6,503 and 5,128, respectively) relative to parental S6P.BA5-1192 (ID50=432) (**Fig. 3b**). This trend in neutralization potency was observed with XBB.1.5 and JN.1 variants although the overall titres were ∼1.5log_10_ lower in comparison to homologous virus. D17 and I1+VI-immune sera possessed geometric mean neutralization ID50s above 70 against XBB.1.5 and JN.1, which emerged after BA.5. By contrast, sera elicited by parental trimer lacked neutralizing activity against XBB.1.5 and JN.1.

**Figure 3.**
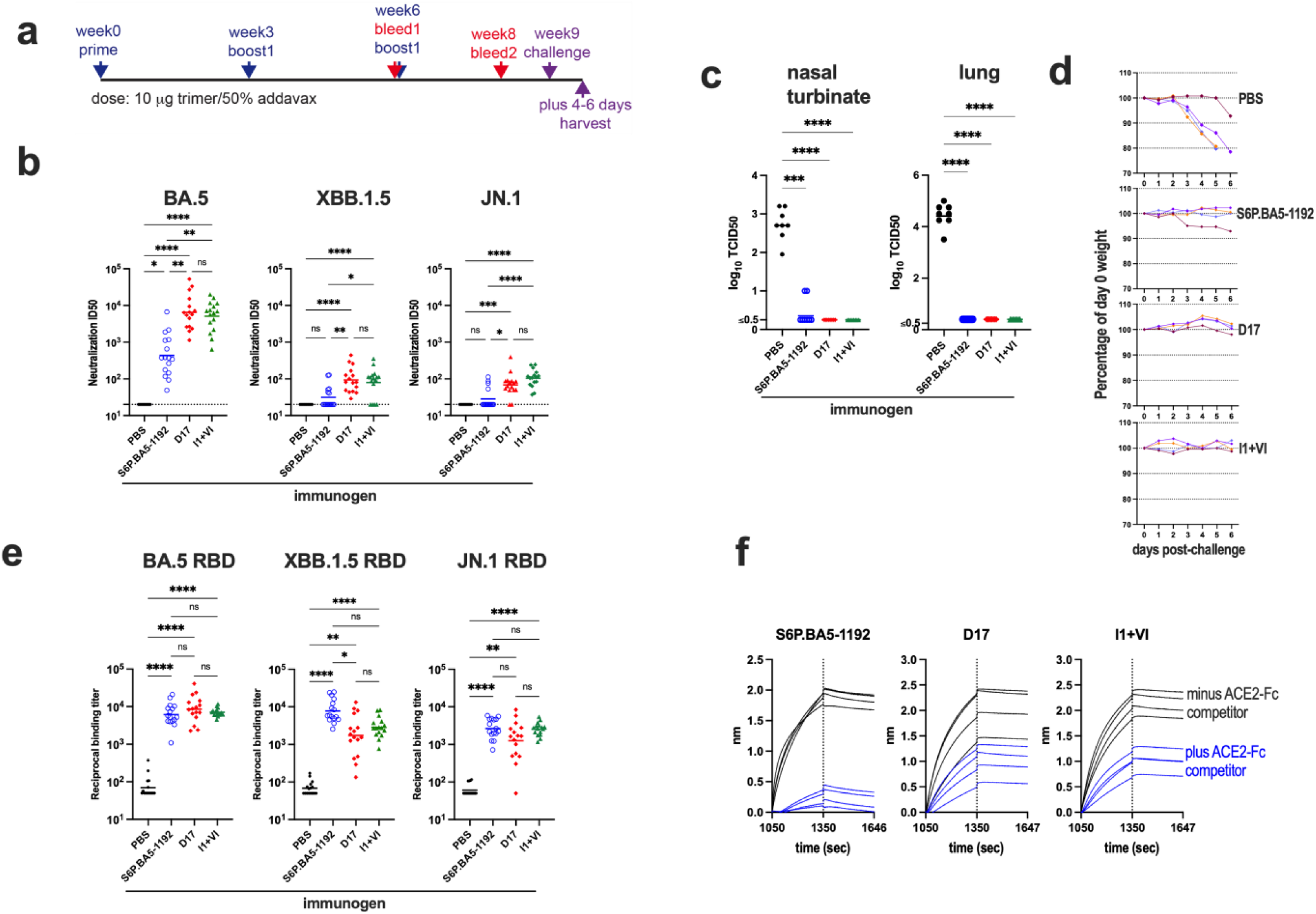
Immunogenicity of soluble covalently stabilized clamp-free omicron BA.5 S trimers. **a.** K18hACE2 mouse immunization-virus challenge protocol. **b.** Neutralization activity of sera obtained after 3 immunizations with S6P.BA5-1192 spike variants. The SARS-CoV-2 variants used in the neutralization assays are shown above the graphs. Horizontal bars are the geometric mean ID50s. **c.** Replication of omicron BA.5 virus in immunized mice after challenge. Virus titres expressed as log_10_ TCID50/mL for NTs and log_10_ TCID50/organ for lungs. Horizontal bars represent geometric mean titres. The lower limit of detection is 10^0.5^ TCID50 per mL for the NTs and 10^0.8^ TCID50 per organ for lungs. **d.** Weight-loss in challenged mice (n=4) expressed as a percentage of day-0 weight. Arrow: mouse euthanized at day 4. **e.** RBD binding titres of sera obtained after 3 immunizations with experimental vaccines. The variant RBDs are indicated above the graphs. The horizontal bars are the geometric means. **f.** ACE2-Fc blockade of vaccinal serum antibody binding to biotin-omicron BA.5 RBD attached to streptavidin biosensors in BLI. The immunogen groups are indicated above the sensograms. (n=4). For **b, c** and **e**: ns, not significant; *, P < 0.05; **, P < 0.01: ***, P < 0.001; ****, P < 0.0001, Kruskal-Wallis test. A Dunn’s post-test was included for **c**.

Three weeks after dose 3, 12 mice/group were challenged with 10^4^ TCID50 of omicron BA.5 virus. Viral titres in nasal turbinates (NTs) and lungs from 8 mice/group were determined by the Reed and Muench method 4 days later. The vehicle control group had mean virus titres of 10^2.7^ TCID50/ml in NTs and 10^4.4^ TCID50/organ in lungs whereas virus was not recovered from mice receiving the 3 vaccines (limit of detection: 10^0.5^ TCID50) except for 2 mice in the parental spike group from which ∼10 TCID50 was recovered from NTs, suggesting break-through infection (**Fig. 3c**). Weight loss was monitored in the remaining 4 mice/group over 6 days. Four of 4 mice receiving vehicle and 1 of 4 mice receiving parental vaccine exhibited significant weight loss over the 6 days of observation, whereas animals receiving D17 and I1+VI vaccines retained their day-0 weights (**Fig. 3d**). The covalently stabilized S6P.BA5-1192 immunogens elicit protective responses in K18hACE2 mice.

Chemiluminescent immunosorbent assays (CLIA) employing isolated spike domains were used to assess the specificity of vaccinal antibody responses. The 3 immunogens elicited similar RBD-binding titres in the range 1/10^3^-1/10^4^ **(Fig. 3e)**. Interestingly, D17 elicited ∼3-4-fold lower NTD-binding titres relative to the parental immunogen (**Supplementary** Fig. 9d) (a similar trend was observed with I1+VI) suggesting that the NTD response was suppressed by the covalent linkages. Although the D17 and I1+VI trimers exhibited stable interactions with stem-directed mNAbs in BLI, there were no significant increases in stem binding titres in the D17 and I1+VI groups when compared with the parental immunogen group (**Supplementary** Fig. 9d).

The observation that D17 and I1+VI seem to promote RBM-occluded conformations (**Supplementary** Fig. 5,6**,7**) prompted the use of an ACE2 competition assay in BLI to gauge the presence of antibodies directed to the RBM region in vaccinal sera. Biotin-BA.5 RBD was attached to streptavidin biosensors and then saturated with ACE2-Fc. After washing, sera were applied; the resulting sensograms are shown in **Fig. 3f**. ACE2-Fc effected almost complete blockade of parental S6P.BA5-1192-elicited sera indicating focussed responses to epitopes involving the RBM. By contrast, D17- and I1+VI-elicited sera were only partially blocked by ACE2-Fc consistent with lower levels of RBM-directed antibodies and the presence of antibodies directed to other parts of the RBD. In **Supplementary** Fig. 9c, ACE2-Fc and mNAbs were used as competitors to examine the footprint of the ACE2 competition assay. As expected, ACE2-Fc completely blocked the binding of ACE2-Fc to the sensor-bound RBD and effected partial inhibition of mNAbs directed to RBM-overlap epitopes (Omi-18, Omi-42 and SA55). By contrast, ACE2-Fc lacked inhibitory activity against broadly neutralizing mNAbs directed to conserved epitopes on the flanks of the RBD (S2H97, SP1-77 and S309). The data suggest that the RBM-occluded nature of D17 and I1+VI elicits broad NAb responses directed to conserved regions of the RBD in addition to the RBM, whereas the NAb response to parental S6P.BA5-1192 is focussed on the RBM, which may explain in part the very low neutralization potency against the emergent isolates, XBB.1.5 and JN.1.

### Covalent stabilization of soluble S oligomers from an emergent SARS CoV-2 variant and divergent ACE2-using bat sarbecoviruses

The Sarbecovirus subgenus comprises 3 ACE2-using clades ^6^: clade 1b, which includes SARS CoV-2 and related Asian bat and pangolin viruses; clade 1a, which includes SARS CoV and related Asian bat and civet viruses; and clade 3, which includes viruses from European and African bats (**Fig. 4a).** An alignment of spike sequences from a sample of isolates identified A570-, D571- and S967-homologous amino acids for D17 and I1 mutagenesis (**Supplementary** Fig. 10a) that were validated in a 3-dimensional context in AlphaFold 3 models ^27^ (**Supplementary** Fig. 10b). D17 and I1+VI were introduced to S6P-1192 constructs derived from an emergent SARS CoV-2 variant (omicron BA.2.86) and examples of divergent ACE2-using bat Sarbecoviruses with potential for spill-over into humans (WIV1 and PRD-0038) ^5,28-30^. Omicron BA.2.86, WIV1 and PRD-0038 share 97.3%, 76.7% and 72.6% amino acid identity, respectively, with the omicron BA.5 spike.

**Fig. 4.**
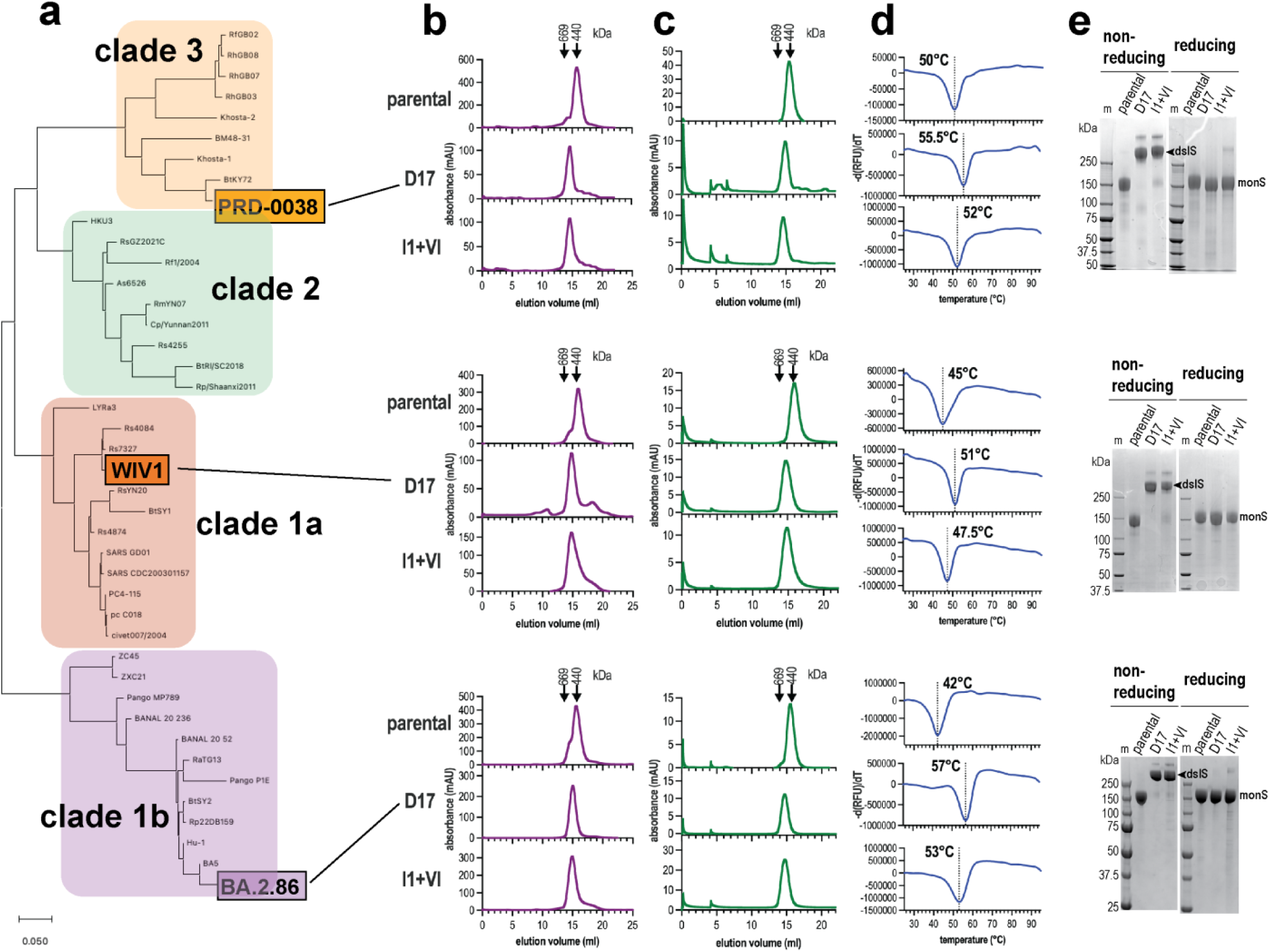
Soluble covalently stabilized clamp-free S trimers from divergent sarbecoviruses. **a.** Maximum likelihood phylogenetic tree of Sarbecovirus spike amino acid sequences. The tree was constructed using MEGA X. **b.** Superose 6 SEC of secreted spike variants eluted from TALON resin. **c.** SEC of purified spike glycoproteins from **b** following a freeze (-80°C)-thaw cycle. **d.** Melting temperature of purified S oligomers determined by DSF. **e.** SDS-PAGE of purified Spike oligomers. dslS, disulfide-linked spike; monS; spike monomer.

Superose 6 SEC of TALON-purified parental spikes revealed major putative trimer peaks eluting close to the 440 kDa standard, while the D17- and I1+VI-mutant glycoproteins eluted earlier, again consistent with larger hydrodynamic radii for the latter (**Fig. 4b**). The 9 glycoproteins were purified to homogeneity (**Fig. 4c**). DSF indicated Tms of 50°C, 45°C and 42°C for parental PRD-0038, WIV1 and BA.2.86 S6P-1192 glycoproteins, respectively, and their Tms were substantially increased by the D17 and I1+VI mutations (**Fig. 4d**). Nonreducing SDS-PAGE indicated that >95% of the purified D17- and I1+VI oligomers were covalently stabilized (**Fig. 4e**). The 1192 stem truncation in concert with the D17 and I1+VI mutations can thus be used to produce covalently stabilized clamp-free soluble spike trimers from divergent sarbecoviruses with increased thermal stability.

BLI was used to assess the epitope profiles of the Sarbecovirus S6P-1192 constructs with pan-Sarbecovirus neutralizing ligands (**Supplementary** Fig. 11). D17 and I1+VI in BA.2.86 and PRD-0038 S6P-1192 spikes led to diminished ACE2-Fc and S2X259 binding responses (the latter is directed to a cryptic RBD epitope ^31^) suggesting some occlusion of the RBM region. As observed with S6P.BA5-1192, D17 led to more marked decreases in binding response relative to I1+VI. By contrast, the overall binding responses of the WIV1-D17 and -I1+VI spikes to ACE2-Fc and S2X259 were similar to parental, but the substantial off-rates observed with the parental spike were largely negated by D17 and I1+VI. The conserved stem region of the glycoprotein variants was probed with S2P6 ^32^. In all cases, the substantial off-rates observed with parental spikes were negated by D17 and I1+VI corresponding to ∼6-500-fold increases in affinities. The covalently stabilized D17 and I1+VI spikes exhibit stable stem-mNAb interactions. The D17 and I1+VI mutations produce intrinsically stable, clamp-free Sarbecovirus spike trimers.

### Broadly neutralizing antibodies to Sarbecoviruses elicited by a trivalent formulation of covalently stabilized Sarbecovirus spike oligomers

Four groups of 10 C57BL/6 mice received 3 doses of vehicle (PBS), 10 μg of monovalent parental or D17-mutated S6P.BA286-1192 oligomers, or a trivalent formulation comprising 4 μg each of D17-mutated BA.2.86, WIV1 and PRD-0038 S6P-1192 oligomers (D17-Trivalent) in AddaVax adjuvant. The mice were bled 2 weeks after the 3^rd^ dose (**Fig. 5a**). CLIA indicated that monovalent and D17-Trivalent immunogens elicited potent cross-reactive binding antibodies to S6P-1192 trimers derived from the 3 isolates (**Fig. 5b**). Potent cross-reactive binding responses to the isolated RBDs were also observed, however the D17-Trivalent immunogen elicited ∼10-fold higher geometric mean binding titres to the WIV1 and PRD-0038 RBDs relative to the monovalent BA.2.86 vaccines (**Fig. 5c**). Potent binding responses were also observed against the BA.2.86 NTD and ancestral stem region for all 3 immunogens (**Fig. 5d,e)**.

**Figure 5.**
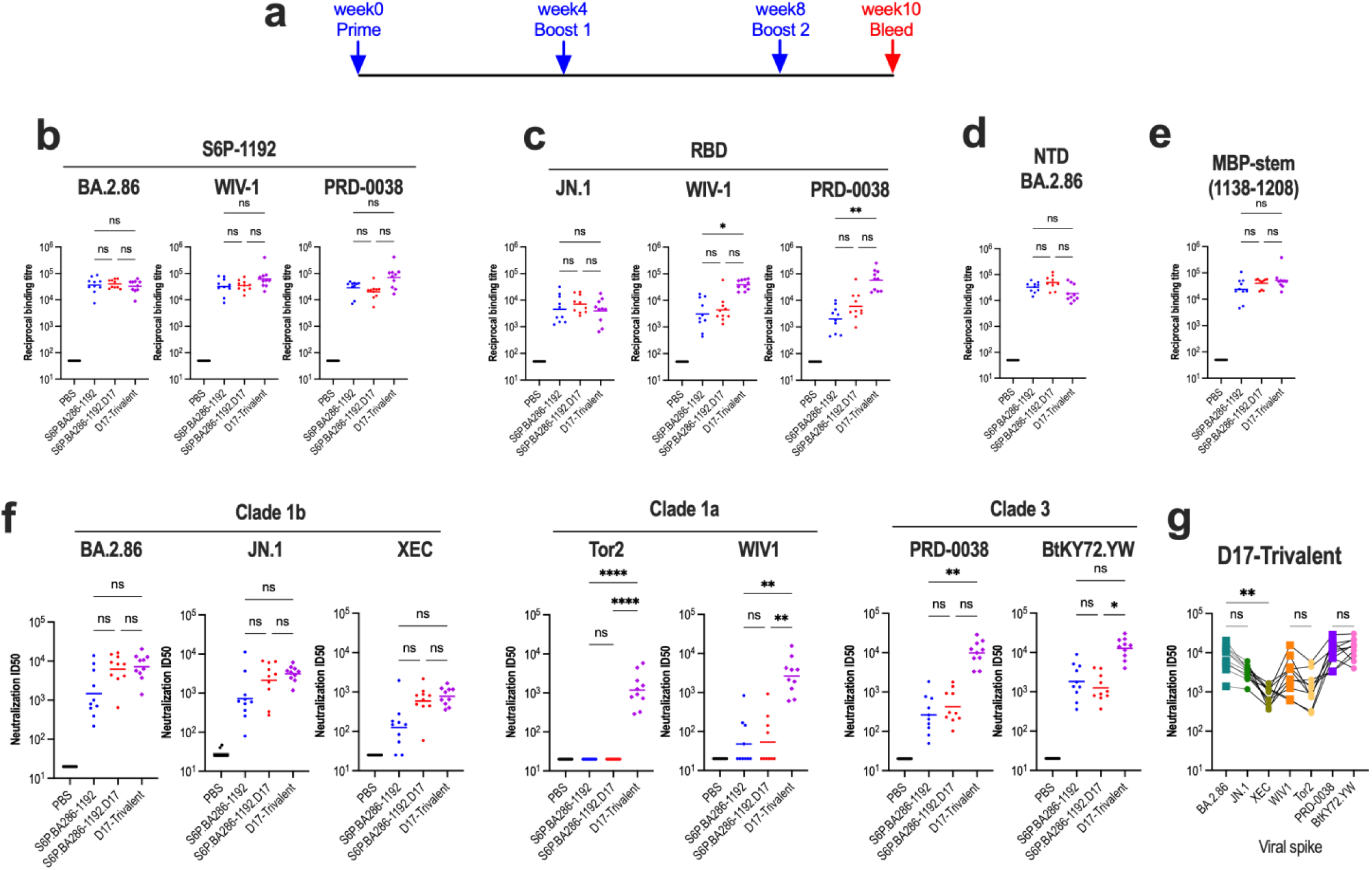
Immunogenicity of a trivalent vaccine comprising covalently stabilized clamp-free S trimers from divergent sarbecoviruses. **a.** C57BL/6 mouse immunization protocol. **b.** S6P-1192 trimer binding titres of vaccinal sera. The variant spikes are indicated above the graphs. **c.** RBD binding titres of vaccinal sera. The variant RBDs are indicated above the graphs. **d.** BA.2.86 NTD binding titres of sera. **e.** MBP-stem(1138-1208) binding titres of sera. **f.** Neutralization activity of sera. The Sarbecovirus variants used in the neutralization assays are shown above the graphs. **g.** Neutralization activity in trivalent spike-immune sera linked by individual mouse. ns, not significant; *, P < 0.05; **, P < 0.01; ****, P < 0.0001, Kruskal-Wallis test; Friedman test for panel **g**. Statistics for PBS group *versus* antigen groups: ns for S6P.BA286-1192 group with XEC and PRD-0038, and S6P.BA286-1192 and S6P.BA286-1192.D17 groups with Tor2 and WIV1; all other antigen groups: P < 0.01.

The D17-mutated monovalent and D17-Trivalent immunogens elicited 3-5-fold higher geometric mean neutralization ID50s against SARS CoV-2 omicron BA.2.86, JN.1 and XEC viruses relative to the monovalent parental immunogen (**Fig. 5f**), a trend that is consistent with that seen with S6P.BA5-1192 parental versus D17 immunogens (**Fig. 3b**). The D17-Trivalent immunogen elicited potent crossneutralizing responses against BA.2.86 JN.1 and XEC viruses, and to clade 1a (WIV1 and SARS CoV Tor2) and clade 3 (PRD-0038 and BtKY72.YW) S-HIV pseudotypes (geometric mean ID50 range 780-12,752) (**Fig. 5f,g**). Significant cross-neutralization of clade 3 spike pseudotypes was observed with monovalent clade 1b spike-elicited immune sera but at a lower level than D17-Trivalent sera. The data indicate that the D17-Trivalent formulation elicits broad and potent neutralizing antibody responses.

Strong correlations between BA.2.86, JN.1, PRD0038 and BtKY72.YW neutralization ID50 with clade-matched S6P-1192 trimer binding titres were observed for D17-Trivalent sera, while moderate correlations with MBP-stem(1138-1208) binding titre were suggested. By contrast, correlations between RBD titre and ID50 were not obvious **(Supplementary** Fig. 12). The data suggest that neutralization in these cases involves a broad NAb response to the S trimer that includes the conserved stem and is not focussed on the RBD. For the other cases, XEC, WIV1 and SARS CoV Tor2, clear correlations were not observed indicative of a yet to be defined neutralization mechanism.

### Covalent stabilization improves the long-term stability of trimeric S vaccines

The stability of parental, D17 and I1+VI S6P.BA286-1192 oligomer samples following incubation for extended periods at 37°C was assessed by SEC, DSF, SDS-PAGE and BLI. SEC of parental S6P.BA286-1192 revealed an additional higher molecular weight species that appeared after 7 days at 37°C and increased in amount over the 112 days of incubation at 37°C **(Fig. 6a)**. The novel higher mol.wt form may correspond to a species with a Tm of 63°C following the heat treatment **(Fig. 6b)**. By contrast, D17 and I1+VI retained their oligomeric structures and Tms following the 112-day incubation. The binding response of parental S6P.BA286-1192 to RBD (ACE2-Fc and S2H97) and stem (S2P6) ligands diminished with treatment at 37°C **(Fig. 6c)**, whereas D17 and I1+VI retained binding activity to S2H97 and S2P6 for the 112 days. As observed with their S6P.BA5-1192 counterparts, D17 and I1+VI exhibited reduced binding to ACE2-Fc, indicating occluded RBMs even with up to 112 days of heat the treatment **(Fig. 6c)**. SDS-PAGE revealed that the heat-treated samples had almost identical migration patterns to their untreated counterparts under non-reducing conditions with a low level of proteolytic clipping revealed under reducing conditions **(Fig. 6c)**. The data indicate that D17 and I1+VI stabilize S6P.BA286-1192 trimers against structural disruption following extended periods at body temperature.

**Fig. 6.**
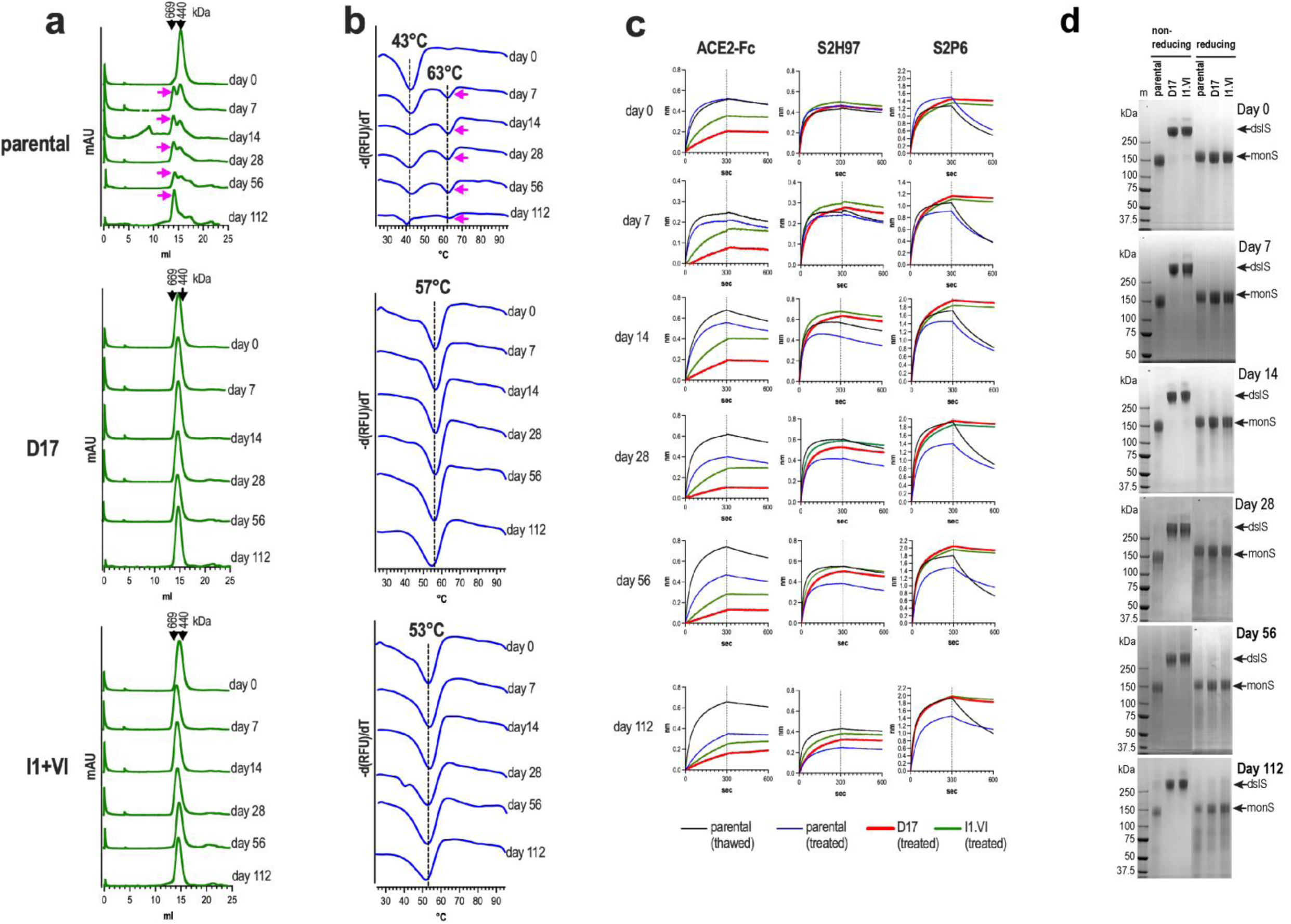
Stability of soluble covalently stabilized clamp-free omicron BA.2.86 S oligomers. **a.** Superose 6 SEC. **b.** DSF. The magenta arrows indicate novel glycoprotein species that appear after 7 days of treatment. **c.** Reactivity with ACE2-Fc and mNAbs in BLI. **d.** SDS-PAGE under non-reducing and reducing conditions. The gels were stained with Coomassie brilliant blue dye. dslS, disulfide linked S; monS, S monomers.

In parallel, parental, D17 and I1+VI S6P.BA286-1192 oligomers were lyophilized and stored at room temperature. The lyophilized samples were periodically reconstituted in sterile deionised water and their biophysical and antigenic integrity assessed as described above. The SEC, DSF, BLI and SDS-PAGE profiles of the reconstituted parental and D17 spikes were virtually identical to their freshly thawed counterparts whereas some evidence of aggregation was observed for I1+VI at the 6-month time point **(Supplementary** Fig. 13). The data indicate that the S6P.BA286-1192 spikes can be stored in lyophilized form for an extended period.

### Expression of covalently stabilized soluble and membrane-bound S6P.BA286-1192 forms from mRNA

We asked whether mRNA could also be used to produce clamp-free covalently stabilized spike trimers. mRNAs encoding parental S6P.BA286-1192, and D17 and I1+VI mutants thereof were transfected into Expi293F cells and expressed at 37°C for 2 days. Spike proteins present in culture supernatants were purified as above. SEC revealed prominent peaks eluting close to the 440 kDa marker (**Supplementary** Fig. 14a) and DSF indicated Tms of 43°C for parental S6P.BA286-1192 and 57.5°C and 55°C for the D17 and I1+VI versions (**Supplementary** Fig. 14b), respectively. These Tms correspond well to those observed for their DNA-expressed counterparts (**see Fig. 4d**). SDS-PAGE under non-reducing conditions confirmed quantitative disulfide stabilization of spike oligomers with D17 and I1 mutations (**Supplementary** Fig. 14c). In BLI, mRNA-derived S6P.BA286-1192.D17 and I1+VI exhibited diminished binding responses to ACE2-Fc and the RBM-directed mNAb Omi-42 compared to parental, whereas the 3 constructs bound potently to S2H97. Again, D17 and I1+VI stabilized the interaction with the stem-directed mNAb CV3-25. The antigenic architectures of mRNA- and DNA-derived S6P.BA286-1192 spikes are thus comparable with D17 and I1+VI leading to RBM occlusion within trimers.

Current SARS CoV-2 mRNA vaccines deliver the prefusion stabilized spike in full-length membrane anchored form. We therefore examined the D17 and I1+VI mutations in this context. 293T cells were transiently transfected with mRNA or DNA encoding parental, D17 and I1+VI versions of full-length S6P.BA286-1273 and the cell lysates analysed by Western blot. **Supplementary** Fig. 14e shows 2 high molecular weight bands (>250 kDa) under non-reducing conditions for D17 and I1+VI constructs that are resolved to the S monomeric molecular weight (∼170 kDa) when reduced. The dslS*a* and dslb bands are consistent with 3 and 2 covalently linked monomers, respectively, per trimer. Flow cytometry and neutralizing ligands were used to probe the antigenic profile of the 3 mRNA-derived spike variants on the surface of intact cells. The data (**Supplementary** Fig. 14f) show strong staining by ligands directed to the NTD (C1520), RBM (ACE2-Fc, SA55, Omi-18, Omi-42), the RBD flanks (SP1-77, S2H97, S309) and stem (CV3-25, CC95-108 and CC99-103). Interestingly, D17- and I1-mediated occlusion of the RBM was not as evident in this context, suggesting that the RBDs are more mobile in the membrane-bound spike.

## Discussion

C-terminal truncation of the stem coupled with rationally designed intermonomer disulfide bonds linking SD1 of S1 to HR1 of S2 enabled the production of soluble, high-yielding and thermostable prefusion-stabilized Sarbecovirus spike trimers that did not require an exogenous trimerization clamp to maintain quaternary structure. In mice, D17 and I1+VI S6P.BA5-1192 trimers elicited substantially more potent NAb responses than their thermolabile parental counterpart. The more potent responses may be due in part to the RBM occluded nature of the covalently stabilized trimers. The closed RBD state may be desirable from a vaccine perspective because the immunodominance of RBM-associated epitopes, which are under antibody selection and serve as immune evasion hot spots ^33-35^, will be suppressed, potentially favouring antibody responses to more conserved regions of the RBD, as was indicated by the competition data presented in Figure 3F. Antibodies directed to conserved RBD epitopes may account for the SARS CoV-2 cross-neutralizing activity observed in D17 and I1+VI sera in Figs. 3b and 5f. The high stability of D17 and I1+VI trimers at 37°C could also contribute to their enhanced immunogenicity. Maintenance of these closed quaternary structures for extended periods at body temperature may allow for the sustained presentation of appropriate epitopes for B cell activation and antibody maturation towards neutralization potency and breadth ^36,37^. The D17 and I1+VI trimers were also associated with suppressed responses to the NTD, which is also an immunodominant domain and hotspot for NAb selection and escape ^38,39^. D17 and I1+VI-immunized K18hACE2 mice were protected against homologous viral challenge whereas break-through infections were detected in 2 of 8 mice and weight loss in 1 of 4 mice that received the parental immunogen. Covalently stabilization of soluble spike trimers and a closed RBD organization appear to be beneficial for protective immune responses.

Amino acid alignments and structural modelling accurately predicted D17 and I1+VI mutagenesis sites in the omicron BA.2.86 variant and divergent ACE2-using bat Sarbecovirus spikes. Quantitative covalent stabilization of spike trimers with substantial increases in Tms were observed in the 3 spikes tested, suggesting that this is a general method for stabilizing clamp-free soluble Sarbecovirus spike trimers. A trivalent formulation comprising S6P-1192.D17 homologs derived from the 3 ACE2-utilizing Sarbecovirus clades (D17-Trivalent) elicited potent and broad cross-clade NAb responses. In addition to potent neutralization of viruses carrying vaccine-homologous spikes, potent neutralization of heterologous spikes derived from the same clades was also observed. For clade 1b and 3 viruses, neutralization ID50 largely correlated with S6P-1192 trimer- and stem-binding titres suggesting a broad NAb response to the S trimer that includes the conserved stem. This may be related, in part, to the D17- and I1+VI-induced stabilization of the interaction between the stem and mNAbs with broad Sarbecovirus neutralizing properties ^32,40^ observed in BLI. High avidity and sustained interactions between the stem of D17- and I1+VI-stabilized trimers and CV3-25-, S2P6-related B cell receptors could result in more potent NAbs to this conserved region. The data suggest that allostery operates between the envelope-distal head and envelope-proximal stem regions of S. An ∼10-fold reduction in neutralization ID50 was observed for SARS CoV-2 XEC relative to the BA.2.86 vaccine strain. Compared to BA.2.86, XEC has 2 and 3 mutations in the NTD and RBD, respectively, suggesting that a component of the D17-Trivalent NAb response is directed to these variable domains where NAb evasion is known to occur ^41^.

Multivalent RBD nanoparticle vaccines in which multiple RBD copies derived from SARS CoV-2 or a variety of sarbecoviruses are assembled on engineered protein scaffolds have been shown to elicit potent and broad NAb responses in animal models ^42-45^. The clamp-free trimeric spikes described here represent an alternative approach for eliciting broad pan-Sarbecovirus NAb responses that target a broad array of epitopes that also include the RBD.

The advent of modified mRNA-lipid nanoparticle technologies has enabled the development and deployment of highly effective vaccines for SARS CoV-2 in record time. The approach used here to produce thermostable clamp-free Sarbecovirus soluble spike trimers was readily adaptable to the mRNA platform with parental, D17 and I1+VI versions of S6P.BA286-1192 expressed from mRNA exhibiting virtually identical biophysical and antigenic properties to their counterparts expressed from DNA. Furthermore, covalent stabilization of full-length membrane-anchor-containing S6P glycoproteins was also achieved following introduction of the D17 and I1+VI mutations illustrating the utility of these changes in other vaccine platforms.

In summary, we have described an approach for producing intrinsically stable, soluble Sarbecovirus spike trimers that are free of potentially immunogenic trimerization clamps derived from foreign proteins, can retain their structure and activity at 37°C and after lyophilization and storage for extended periods. We also demonstrate that these immunogens can elicit potent and protective neutralizing antibody responses and can be readily applied to the mRNA platform. The technology has the potential to simplify the manufacture, storage and distribution of trimeric recombinant Sarbecovirus spike protein vaccines and increase the potency of other vaccine modalities such as mRNA to mitigate future pandemics.

## Materials and Methods

### Recombinant spike vectors

S2P.BA5.VI-1208 (previously referred to as S2P.OmiBA45.VI-1208) is described in ^14^. Polymerase chain reaction was used to truncate its stem to S1147, K1157, D1165, N1192, D1199, L1200, Q1201 and G1204. Cysteine substitution, 6P and V1016A/I1020A mutations were introduced to various spike expression vectors using synthetic DNA (GeneART-ThermoFisher Scientific). Synthetic DNA encoding parental, D17 and I1+VI S6P-1192 sequences derived from omicron BA.2.68, PRD-0038 and WIV1 (Genbank accession numbers: OR775659, QTJ30153, AGZ48828, respectively) were obtained from Genscript. The N-terminal domain (amino acids 16-305) and RBD (amino acids 332-532) derived from omicron BA.5, XBB and JN.1 SARS CoV-2 isolates were obtained from Genscript. All spike sequences incorporate an N-terminal tissue plasminogen activator leader sequence linked via Ala-Ser to the first amino acid of the mature spike protein or fragment thereof and Gly-Ser-Gly-Ser-His_6_ at the C-terminus except for RBD, which has Gly-Ser-Gly-Ser-His_8_-Gly-Ser-Gly-Ser-avitag. pcDNA3 (Thermo-Fisher Scientific) was the expression vector. All Spike DNA sequences were codon-optimised for expression in human cells and verified by Sanger sequencing with ABI BigDye Terminator 3. The MBP-stem(1138-1208) chimera comprises *E. coli* MBP linked to the ancestral stem amino acids 1138-1208 via a tri-alanine linker_46._

### Expression and purification of recombinant spike proteins

Spike expression vectors were transfected into Expi293F cells using Expifectamine, as recommended by the manufacturer (ThermoFisher Scientific). The cells were cultured for 7 days at 34°C after which the transfection supernatants were clarified by centrifugation and filtration through 0.45 μm nitrocellulose filters. The glycoproteins were then purified by divalent cation affinity chromatography using TALON resin (Merck) followed by SEC using a Superose 6 Increase 10/300 column linked to an AKTApure instrument (Cytiva). The molecular weight markers were thyroglobulin (669 kDa), ferritin (440 kDa), and aldolase (158 kDa). The buffer was Dulbecco’s phosphate buffered saline pH 7.4. MBP-stem(1138-1208) was induced in BL21(DE3) cells and purified as described ^47^. SDS-PAGE was performed using NuPAGE precast gels and an XCell SureLock Mini-Cell Electrophoresis System (Thermo Fisher). Precision plus prestained protein standards (Bio-Rad) were used.

### Synthetic peptide

A dynthetic peptide corresponding to the fusion peptide (amino acids 808–832, DPSKPSKRSFIEDLLFNKVTLADAG) was synthesized by Genscript with an N-terminal biotin moiety and C-terminal amide.

### Recombinant spike ligands

*<u>hACE2-Fc</u>* comprises amino acids 19-615 of human ACE2 linked to the IgG1 Fc domain via Gly-Ser ^14^. *<u>Recombinant mNAbs.</u>* pCDNA3-based human IgG1 heavy and kappa and lambda light chain expression vectors ^48^ containing the variable regions of SARS CoV-2 directed mNAbs S2H97 ^6^, S2X59 ^31^, Omi-18 and Omi-42 ^17^, S309 ^49^, SA55 ^50^ SP1-77 ^18^, C1520 ^51^, COV44-79 ^19^, CV3-25 ^40^, S2P6 ^32^, CC95-108, CC99-103 ^52^ were produced in-house as described ^48^.

### Thermostability

Differential scanning fluorimetry employing SYPRO orange and a QuantStudio 7 Real-time qPCR System was used to assess protein thermostability ^53^. The Tm was determined as the minimum of the negative first derivative of the melting curve. Alternatively, filter-sterilized purified spike samples were stored at 37°C in a humidified CO_2_ tissue culture incubator after which they were analyzed by Superose 6 SEC, SDS-PAGE, DSF and BLI with ACE2-Fc, S2H97 and S2P6 antibodies.

### Biolayer interferometry

BLI-based measurements were determined using an OctetRED96 System (ForteBio, Fremont CA) as described ^14^. Streptavidin biosensors (ForteBio) were used for epitope binning experiments. The experiments included 6 steps: (a) baseline (180 s); (b) biotin-RBD loading (300 s); (c) second baseline (75 s); (d) binding of ACE2-Fc (650 s), (e) association of test antibody (300 sec), (f) dissociation of test antibody (300 s). Fitting curves were constructed using Octet Analysis Studio 13.0.3.52 software using a 1: 1 binding model, and double reference subtraction was used for correction.

### Mice

Eight-to-10-week-old B6.Cg-Tg(K18-ACE2)2Prlman female mice (Animal Resources Centre, Australia) were inoculated at weeks 0, 3 and 6 via interscapular subcutaneous injection with 10 μg spike vaccines or PBS (vehicle) in 50% v/v Addavax (16 animals per group). A pilot mandibular bleed was taken one day before and 2 weeks after dose 3 (8 animals). Terminal bleeds of 4 mice/group was also performed. Three weeks following dose 3, all remaining mice were lightly anesthetized using 4% v/v isoflurane and infected intranasally with 10^4^ TCID50 of SARS-CoV-2 Omicron BA.5 (hCoV-19/Australia/VIC61194/2022 [B.1.1.529.5] GISAID: EPI_ISL_13276063) (Victorian Infectious Diseases Reference Laboratory) in 50 μL of PBS. Four mice were treated with 50 μL PBS as a mock challenge. At four days post infection 8 mice/group were culled by CO_2_ asphyxiation before lungs and nasal turbinates were homogenized in 2 mL and 1 mL of PBS, respectively. Weights of 4 mice/group were measured daily over 6 days. The experiment was conducted in a certified Physical Containment Level 3 (PC3) facility (AgriBio, Centre for AgriBioscience, Bundoora, Australia), under approval from the University of Melbourne Animal Ethics Committee, ethics number 20763. In a second experiment, 8–10-week-old C57BL/6 female mice (Walter and Eliza Hall Institute of Medical Research) were inoculated at weeks 0, 4 and 8 via interscapular subcutaneous injection with 10 μg monovalent spike vaccines, or 4 μg of each spike in a trivalent formulation (D17-Trivalent), or PBS (vehicle) in 50% v/v Addavax (10 animals per group). Terminal bleeds were performed 2 weeks after the 3^rd^ dose. This experiment was performed under approval from the WEHI Animal Ethics Committee, ethics number 2023.012.

### Virus Quantitation

Vero-TMPRSS2 cells were infected in quadruplicate in 24 well and 96-well plates to determine the tissue culture infective dose (TCID50/ml). Cells were incubated at 37°C and 5% CO2 for 5 days. Following incubation, wells with CPE were recorded and the virus titre (TCID50/mL) for each sample determined by limited dilution using the Reed and Muench method.

### Authentic virus neutralization assay

SARS-CoV-2 variants were isolated from diagnostic respiratory specimens as previously described and through the primary diagnostic provider Douglas Hanly Moir under UNSW ethics IREC5100 ^54,55^. Briefly, specimens were sterile-filtered through 0.22 µm column-filters at 10,000 x*g* and incubated on TASL-19 cells for 72 hours. Supernatants from wells showing cytopathic effect were incubated with TASL-19 cells for 24 to 48 h (or until cytopathic effects had led to loss of >50% of the cell monolayer). The supernatant was cleared by centrifugation (2,000 x*g*, 10 min) and stored at -80°C (passage 2). The neutralizing activity of sera against Omicron BA.5 (hCoV-19/Australia/NSW2715/2020 A.2.2), XBB.1.5 (hCoV-19/Australia/NSW-ICPMR-54023/2024), JN.1 (hCoV-19/Australia/NSW-ICPMR-56406/2024) and XEC (hCoV-19/Australia/NSW-ICPMR-29513/2022) SARS-CoV-2 was determined with the rapid high-content SARS-CoV-2 microneutralization assay ^26^. Briefly, Hoechst-33342-stained HAT-24 cells were seeded in 384-well plates (Corning, CLS3985). Serially diluted heat-inactivated (56°C, 30 min) vaccinal sera were co-incubated with an equal volume of SARS-CoV-2 virus solution at twice the median lethal dose for 1 h at 37 °C. 40 μl of serum-virus mixtures were added to an equal volume of pre-plated cells, incubated for 20 h and then directly imaged on an ImageXpress Pico Automated Cell Imaging System (Molecular Devices). Cellular nuclei counts were obtained with CellReporterXpress Image Acquisition and Analysis software (Molecular Devices), and the percentage of virus neutralization was calculated as described in Aggarwal et al. 2022. The neutralization ID50 was the last consecutive dilution reaching ≥50% neutralization.

### S-HIV pseudovirus neutralizing assay

S-HIV luciferase reporter viruses were produced by cotransfecting 293T cells with pNL4.3LucR-E-^56^ and pcDNA3 containing DNA inserts encoding the S open reading frames from SARS CoV Tor2 (AAP41037.1), WIV1 (AGZ48828), PRD-0038 (QTJ30153) and BtKY72 (APO40579); the BtKY72 spike contains the K482Y/T487W and is called BtKY72.YW. The viruses were harvested 72 h post transfection. Heat inactivated sera (56°C for 30 minutes) were serially diluted in DMF10 and incubated for 1h at 37°C with an equal volume of S-pseudoviruses. Virus-serum mixtures were added to monolayers of HAT24 cells (for SARS CoV Tor2 and WIV1) or HAT 24 cells transfected with a *Rhinolophus alcyone* expression vector (for PRD-0038 and BtKY72.YW) attached to 96 well plates the day prior at 10,000 cells/well. After 3 days, tissue culture fluid was removed and monolayers lysed with cell culture lysis reagent (Promega) and luciferase measured using luciferase substrate (Promega) in a CLARIOstar plate reader (BMG LabTechnologies). The mean percentage entry was calculated as (RLU plasma+virus)/(RLU medium+virus)*100. The percentage entry was plotted against the reciprocal dilution of plasma in Prism v9.3.0 and curves fitted using the sigmoidal 4PL model. The reciprocal dilution of plasma required to prevent 50% virus entry was calculated from the non-linear regression line (ID50).

### Chemiluminescence Immunoassay (ELISA)

Nunc Maxisorp 384-well white plates were coated with 1 μg/ml of S glycoproteins, NTD proteins, RBD proteins, stem proteins or synthetic peptides overnight at 4°C, washed with PBS and blocked with BSA (10mg/ml, PBS) at room temperature for 1 h. The plates were again washed and incubated with serially diluted serum samples or mNAbs for 2 h at room temperature. Antibody binding was detected using horseradish peroxidase-labelled rabbit anti-guinea pig antibody (Dako, Glostrup, Denmark). Light signals were detected using SuperSignal ELISA Pico Chemiluminescent Substrate (ThermoFisher Scientific) and measured immediately for 0.5 seconds using CLARIOstar (BMG Lab Technologies). Relative light units (RLUs) were plotted against the reciprocal dilution in GraphPad Prism 10.1.0 and curves fitted using Specific binding with Hill slope. The binding titre was defined as the reciprocal dilution of serum giving RLUs fifty-times that of background, as defined by binding to BSA.

### S6P.BA286-1192 glycoprotein stability experiments

Purified, trimeric parental, D17 and I1+VI-mutated S6P.BA286-1192 glycoproteins in PBS were adjusted to 1 mg/mL and supplemented to 0.02% w/v sodium azide. The samples were incubated at 37°C for up to 112 days in a humidified tissue culture incubator in the presence of 5% CO2. At various intervals, the samples were assessed by Superose 6 SEC, DSF in the presence of SYPRO orange, reactivity with ACE2-Fc and human mNAbs in BLI, and SDS-PAGE under reducing and non-reducing conditions.

### Lyophilization

Purified S6P.BA286-1192 protein samples in PBS were sterilized using 0.45 μm syringe filters and adjusted to a final concentration of 2 mg/mL. An equal volume of sterile 5% w/v sucrose/PBS solution was then added for a final protein concentration of 1 mg/mL. Sixteen 100 μL aliquots of each protein in screw capped microcentrifuge tubes were frozen overnight at -80°C. The protein samples were lyophilized for 5 h (2-5 Torr) in a vacuum oven (NAPCO 5831; Selby Scientific) connected to a vacuum pump (AVT Services) and freezer compartment (Christ Beta 2-8 LD-plus; John Morris Scientific). The samples were recapped and stored together with desiccant sachets in a sealed box at room temperature in the dark for the specified times. Each sample was reconstituted in 100μL sterile deionised water and clarified by centrifugation at 20,000 xg for 3 min at 4°C before analysis by SEC, DSF, SDS-PAGE and BLI.

### Cryo-EM

For negative stain EM, S6P-BA286-1192.D17 was loaded onto Quantifoil R 1.2/1.3 holey carbon grids that were previously subjected to glow-discharge for 30 s at 30 mA current, before staining with 2% uranyl acetate. Grids were imaged using a Tecnai Spirit G2 TEM (FEI) operating at 120 kV and equipped with a Gatan Alpine camera. Negative stain EM images were collected at a nominal magnification of 24000x with a defocus of 1.5 to 2 μm and a total exposure of 50 e-/Å2. For cryo-EM, S6P BA286-1192 was loaded onto Quantifoil R 1.2/1.3 holey carbon grids that were previously subjected to glow-discharge for 30 s at 30 mA. The grids were then blotted using a FEI Mark IV vitrobot (Blot force of 5 and blot time of 3.5 s) operating at 4°C and 100% relative humidity, before plunge-freezing in liquid ethane. Frozen grids were imaged using a Titan Krios TEM (FEI) operating at 300 kV and equipped with a K3 camera. Movies were recorded in counting mode at a nominal magnification of 105,000x (corresponding to pixel size of 0.82 Å/pixel), with a total dose of 60 e-/Å^2^ and a final dose per frame of 1 e-/Å^2^/frame. EPU’s AFIS mode was employed to significantly speed up data collection.

### Data Processing

Movies were collected and processed in cryoSPARC 4.6.2, along with their respective beam shift values for exposure group assignments. Movies were pre-processed using CryoSPARC patch Motion Correction to reduce motion blur, and Patch CTF estimation for estimating the contrast transfer function. Movies were manually curated based on average defocus, CTF fit resolution, estimated ice thickness and total full-frame motion distance, which yielded a final set of high-quality 10,387 movies. Initially, particles were picked using blob picker, which were subsequently inspected, extracted, and 2D classified to be used as templates in template picker. A final 4,251,982 particles were extracted using a box size of 440 pixels from the template picking and were subjected to further 2D classification. The particle stack was curated with successive rounds of heterogenous refinement (4 classes) and 2 rounds of ab initio reconstructions (3 classes). Non-uniform refinement was used to improve the map quality, followed by rebalancing to reduce the orientation bias (80%, random selection) and a final non-uniform refinement with 327,643 particles. The final map was refined to a nominal GSFSC resolution of 3.88 Å as reported in cryoSPARC but suffers from significant anisotropy (cFAR 0.36) that precluded model building.

### mRNA synthesis and transfection

mRNAs were produced with HiScribe T7 mRNA synthesis kit (NEB, Australia) using linearized DNA produced by PCR amplification. The mRNAs included de novo designed 5’- and 3’-UTRs and polyA_125_ tails. N1-methyl-pseudoUTP instead of UTP was used to produce chemically modified mRNA. CleanCap reagent AG (TriLink) was used to produce Cap1 chemistry at the 5’ terminus.

The mRNA was subject to cellulose purification before use. mRNAs were transfected into Expi293F or 293T cells using lipofectamine MessengerMAX (Thermo Fisher Scientific). The cells were cultured at 37°C for 48 h. Soluble S glycoproteins purified from culture supernatants as described, above. Membrane anchored glycoproteins were were assessed by SDS-PAGE and Western blotting of 293T cell lysates. The blots were probed with guinea pig S glycoprotein immune sera raised in guinea pigs ^14^.

### Flow Cytometry

Transfected 293T cells were detached using versene solution, resuspended in PBS and stained with LIVE/DEAD near-IR dead cell stain (Invitrogen). The cells were washed in ice-cold FACS buffer (5% v/v fetal calf serum in PBS containing 2 mM ethylenediamine tetra acetic acid) and incubated with ACE2-Fc and human mNAbs in FACS buffer for 1 h at room temperature in V-bottom 96-well culture plates. After further washing, the cells were incubated with AlexaFluor 647 goat anti-human (H+L) (Invitrogen) for 30 min at room temperature in the dark. The cells were washed, resuspended and immediately applied to a Canto II flow cytometer. Ten thousand events were captured for each antibody-S6P.BA286-1273 variant combination. FlowJo software was used for data analysis. The viable 293T cell population was first gated by forward scatter and side scatter and single cells analyzed after doublet discrimination.

## Supporting information

Langer et al Supplementary Material

