## Supplementary material for "High-stability, clamp-free soluble Sarbecovirus spike trimers and their potential for pan-Sarbecovirus vaccine development": Langer et al Supplementary Material

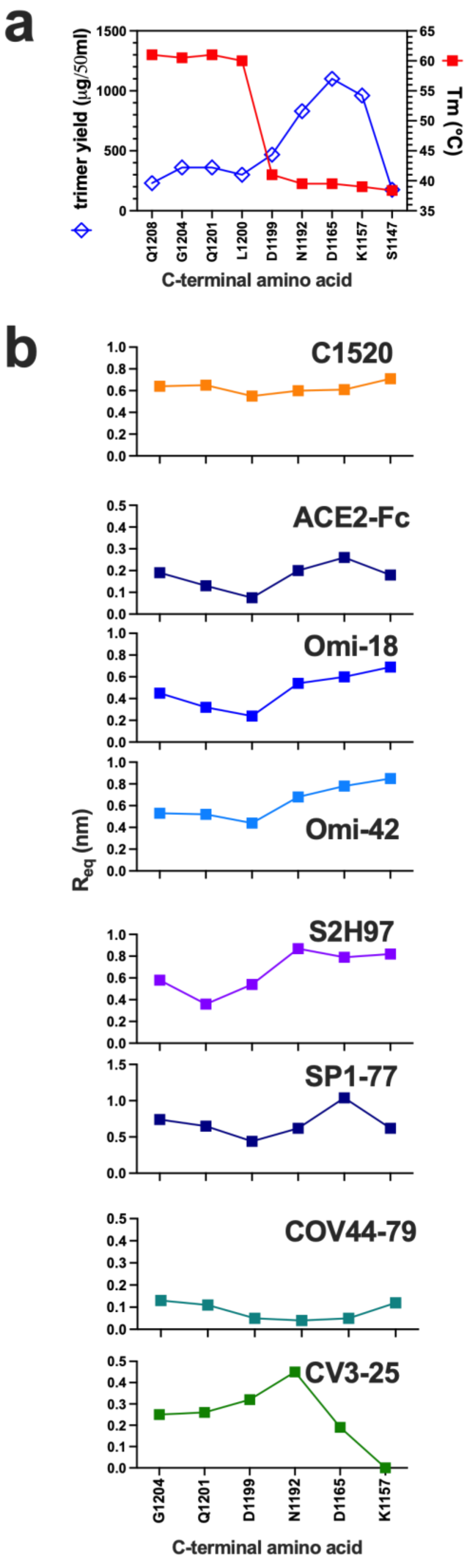


**Supplementary Fig 1a.** Plot of trimer yield and Tm as a function of C-terminal length (data from Fig. 1). **b.** Response (nm) at binding equilibrium (Req) is plotted as a function of C-terminal length (data from Supplementary Table 1).

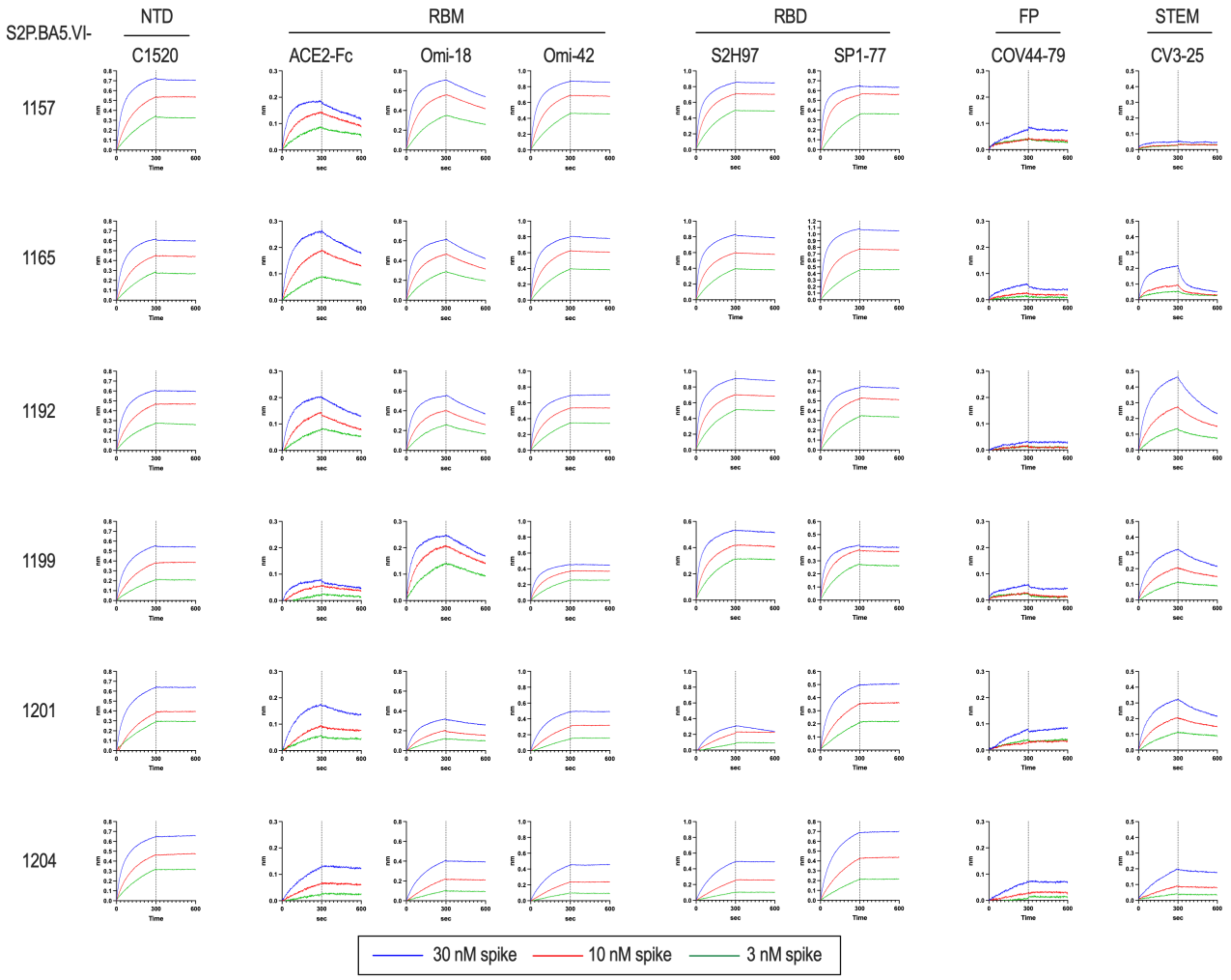


**Supplementary Fig. 2.** Epitope profiles of purified spike oligomers determined in BLI. Neutralizing ligands were immobilized on anti-human IgG Fc capture biosensors and the spike trimers were in the analyte phase. Association phase: 0-300 sec; dissociation phase: 301-600 sec.

**Supplementary Table 1. Binding kinetics of S2P.BA5.VI stem truncation mutants to ACE2-Fc and human monoclonal NAbs.**

| Ligand | C-terminal amino acid | $R_{eq}$[a] (nm) | $R^2$[b] | $\chi^2$[c] | KD (M) | $k_{on}$ (1/Ms) | $k_{dis}$ (1/sec) |
|---|---|---|---|---|---|---|---|
| **C1520** | K1157 | 0.71 | 0.996 | 0.276 | $7.4x10^{-11}$ | $6.2x10^{5}$ | $4.6x10^{-5}$ |
| | D1165 | 0.61 | 0.995 | 0.237 | $1.4x10^{-10}$ | $6.2x10^{5}$ | $8.8x10^{-5}$ |
| | N1192 | 0.60 | 0.993 | 0.351 | $8.2x10^{-11}$ | $5.9x10^{5}$ | $4.8x10^{-5}$ |
| | D1199 | 0.55 | 0.997 | 0.135 | $3.8x10^{-11}$ | $4.5x10^{5}$ | $1.7x10^{-5}$ |
| | Q1201 | 0.65 | 0.996 | 0.217 | $<10^{-12}$ | $4.1x10^{5}$ | $<10^{-7}$ |
| | G1204 | 0.64 | 0.991 | 0.484 | $<10^{-12}$ | $5.4x10^{5}$ | $<10^{-7}$ |
| **ACE2-Fc** | K1157 | 0.18 | 0.996 | 0.015 | $1.9x10^{-9}$ | $7.8x10^{5}$ | $1.5x10^{-3}$ |
| | D1165 | 0.26 | 0.998 | 0.015 | $2.5x10^{-9}$ | $5.0x10^{5}$ | $1.2x10^{-3}$ |
| | N1192 | 0.20 | 0.994 | 0.033 | $3.6x10^{-9}$ | $4.5x10^{5}$ | $1.6x10^{-3}$ |
| | D1199 | 0.07 | 0.970 | 0.027 | $3.1x10^{-9}$ | $4.4x10^{5}$ | $1.4x10^{-3}$ |
| | Q1201 | 0.13 | 0.996 | 0.018 | $3.2x10^{-9}$ | $2.8x10^{5}$ | $8.9x10^{-4}$ |
| | G1204 | 0.19 | 0.997 | 0.009 | $1.5x10^{-9}$ | $1.2x10^{5}$ | $1.8x10^{-4}$ |
| **Omi-18** | K1157 | 0.69 | 0.991 | 0.477 | $1.2x10^{-9}$ | $7.6x10^{5}$ | $9.2x10^{-4}$ |
| | D1165 | 0.60 | 0.992 | 0.312 | $1.7x10^{-9}$ | $7.3x10^{5}$ | $1.3x10^{-3}$ |
| | N1192 | 0.54 | 0.993 | 0.248 | $2.1x10^{-9}$ | $6.8x10^{5}$ | $1.4x10^{-3}$ |
| | D1199 | 0.24 | 0.988 | 0.072 | $1.6x10^{-9}$ | $8.1x10^{5}$ | $1.3x10^{-3}$ |
| | Q1201 | 0.32 | 0.996 | 0.051 | $2.0x10^{-9}$ | $3.8x10^{5}$ | $7.7x10^{-4}$ |
| | G1204 | 0.45 | 0.999 | 0.019 | $4.0x10^{-10}$ | $2.4x10^{5}$ | $9.6x10^{-5}$ |
| **Omi-42** | K1157 | 0.85 | 0.992 | 0.773 | $7.4x10^{-12}$ | $7.6x10^{5}$ | $5.6x10^{-6}$ |
| | D1165 | 0.78 | 0.991 | 0.751 | $5.5x10^{-11}$ | $7.6x10^{5}$ | $4.2x10^{-5}$ |
| | N1192 | 0.68 | 0.990 | 0.627 | $<10^{-12}$ | $6.5x10^{5}$ | $<10^{-7}$ |
| | D1199 | 0.44 | 0.988 | 0.283 | $<10^{-12}$ | $7.7x10^{5}$ | $<10^{-7}$ |
| | Q1201 | 0.52 | 0.998 | 0.084 | $<10^{-12}$ | $3.0x10^{5}$ | $<10^{-7}$ |
| | G1204 | 0.53 | 0.999 | 0.016 | $<10^{-12}$ | $2.1x10^{5}$ | $<10^{-7}$ |

| Ligand | C-terminal amino acid | $R_{eq}$ (nm) | $R^2$ | $\chi^2$ | KD (M) | $k_{on}$ (1/Ms) | $k_{dis}$ (1/sec) |
|---|---|---|---|---|---|---|---|
| **S2H97** | K1157 | 0.82 | 0.979 | 1.634 | $<10^{-12}$ | $1.0x10^{6}$ | $<10^{-7}$ |
| | D1165 | 0.79 | 0.989 | 0.952 | $4.1x10^{-11}$ | $9.4x10^{5}$ | $3.8x10^{-5}$ |
| | N1192 | 0.87 | 0.981 | 1.614 | $1.3x10^{-11}$ | $9.7x10^{5}$ | $1.2x10^{-5}$ |
| | D1199 | 0.54 | 0.981 | 0.538 | $2.4x10^{-11}$ | $9.9x10^{5}$ | $2.4x10^{-5}$ |
| | Q1201 | 0.36 | 0.986 | 0.230 | $1.63x10^{-9}$ | $2.2x10^{5}$ | $3.5x10^{-4}$ |
| | G1204 | 0.58 | 0.999 | 0.015 | $<10^{-12}$ | $2.1x10^{5}$ | $<10^{-7}$ |
| **SP1-77** | K1157 | 0.62 | 0.993 | 0.373 | $<10^{-12}$ | $1.2x10^{6}$ | $<10^{-7}$ |
| | D1165 | 1.04 | 0.996 | 0.707 | $<10^{-12}$ | $9.3x10^{5}$ | $<10^{-7}$ |
| | N1192 | 0.62 | 0.989 | 0.531 | $<10^{-12}$ | $8.1x10^{5}$ | $<10^{-7}$ |
| | D1199 | 0.44 | 0.983 | 0.300 | $8.5x10^{-11}$ | $1.1x10^{6}$ | $9.2x10^{-5}$ |
| | Q1201 | 0.65 | 0.997 | 0.119 | $<10^{-12}$ | $4.4x10^{5}$ | $<10^{-7}$ |
| | G1204 | 0.74 | 0.998 | 0.120 | $<10^{-12}$ | $3.1x10^{5}$ | $<10^{-7}$ |
| **COV44-79** | K1157 | 0.12 | 0.913 | 0.058 | $6.7x10^{-9}$ | $1.1x10^{5}$ | $7.2x10^{-4}$ |
| | D1165 | 0.05 | 0.967 | 0.014 | $3.8x10^{-9}$ | $3.81x10^{5}$ | $1.4x10^{-3}$ |
| | N1192 | 0.04 | 0.934 | 0.010 | $2.7x10^{-9}$ | $1.5x10^{5}$ | $4.2x10^{-4}$ |
| | D1199 | 0.05 | 0.858 | 0.050 | $2.7x10^{-9}$ | $5.5x10^{5}$ | $1.5x10^{-3}$ |
| | Q1201 | 0.11 | 0.969 | 0.028 | $<10^{-12}$ | $1.2x10^{5}$ | $<10^{-7}$ |
| | G1204 | 0.13 | 0.980 | 0.023 | $<10^{-12}$ | $8.5x10^{4}$ | $<10^{-7}$ |
| **CV3-25** | K1157 | 0.05 | 0.915 | 0.019 | na | na | na |
| | D1165 | 0.19 | 0.965 | 0.186 | $1.1x10^{-8}$ | $2.6x10^{5}$ | $6.1x10^{-3}$ |
| | N1192 | 0.45 | 0.997 | 0.072 | $5.8x10^{-9}$ | $4.0x10^{5}$ | $2.3x10^{-3}$ |
| | D1199 | 0.32 | 0.995 | 0.062 | $3.1x10^{-9}$ | $4.1x10^{5}$ | $1.25x10^{-3}$ |
| | Q1201 | 0.26 | 0.997 | 0.025 | $3.4x10^{-9}$ | $2.3x10^{5}$ | $7.7x10^{-4}$ |
| | G1204 | 0.25 | 0.996 | 0.025 | $2.4x10^{-9}$ | $1.6x10^{5}$ | $3.8x10^{-4}$ |

[a]$R_{eq}$: Response units at binding equilibrium;
[b]$R^2$: square of the coefficient of correlation between sensogram data and 1:1 bimolecular binding model;
[c]$\chi^2$:Sum of the squared deviations: sensogram data versus 1:1 bimolecular binding model

### Supplementary Material

**a**

| code | mutation | Cα-Cα distance (Å) | subunits |
|---|---|---|---|
| A2 | T547C/N978C | 6.7, 6.8, 6.4 | S1-S2 |
| B8 | A713C/Q895C | 5.6, 5.7, 5.6 | S2-S2 |
| C12 | A668C/T866C | 5.4, 5.4, 5.5 | S1-S2 |
| D17 | D571C/S967C | 6.9, 5.8, 5.5 | S1-S2 |
| E22 | G757C/S968C | 5.5, 5.1, 4.7 | S2-S2 |
| F24 | S758C/T961C | 7.5, 7.6, 7.7 | S2-S2 |
| G1 | N317C/D737C | 7.5, 7.2, 8.0 | S1-S2 |
| I1 | A570C/S967C | 7.0, 5.5, 5.0 | S1-S2 |
| K19 | A890C/G1046 | 4.7, 3.8, 4.7 | S2-S2 |
| L23 | N914C/S1123C | 7.0, 7.3, 7.4 | S2-S2 |
| M2 | S1030C/D1041 | 5.8, 5.7, 5.8 | S2-S2 |

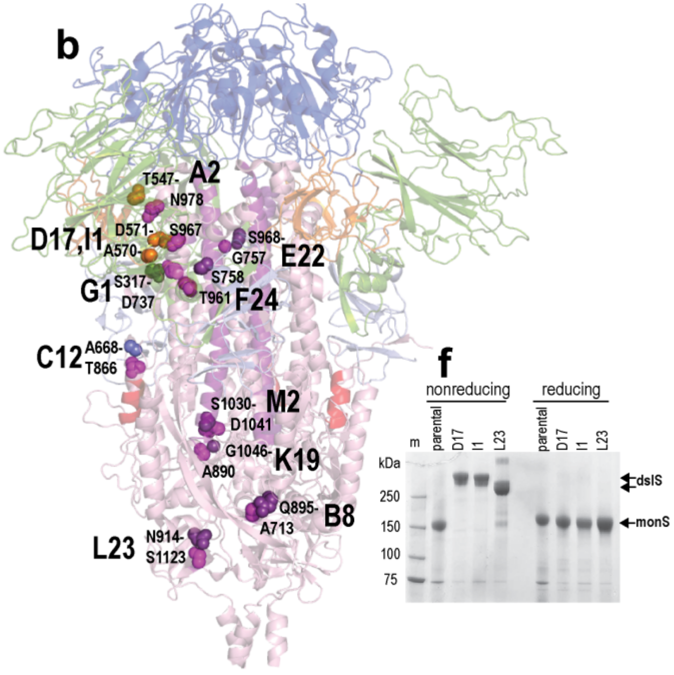


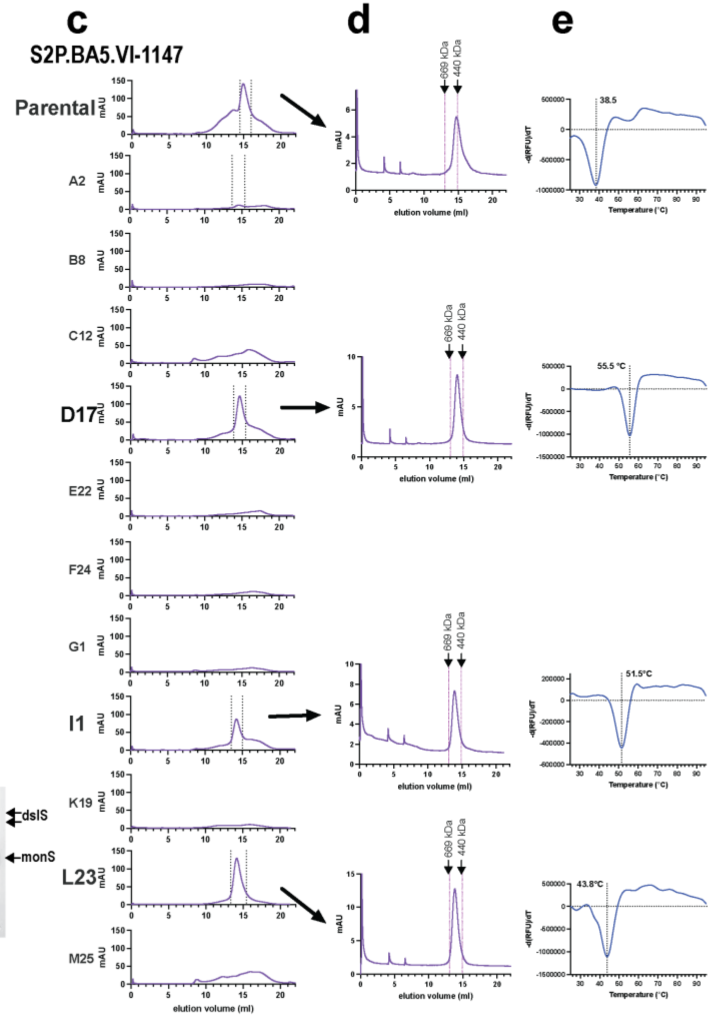


**Supplementary Fig. 3. a.** Near-neighbour contact amino acids targeted with Cys mutations. Contacting residues were identified using the Ligand-Protein Contacts & Contacts of Structural Units server and PDB ID: 6VSB. **b.** Location of paired Cys substitution targets in the omicron BA.5 S6P-foldon trimer. Drawn with PYMOL and PDB ID 7XNQ. Amino acid pairs replaced with Cys are shown in CPK and identified by the code used in A. **c.** SEC of secreted S2P.BA5.VI-1147 spike variants eluted from TALON resin. The fractions within the dotted vertical lines in **C** were pooled, concentrated and then subjected to SEC (**d**) and DSF (**e**) to obtain Tms. **f**. SDS-PAGE of purified spike oligomers. dslS, disulfide-linked spike; monS; spike monomer.

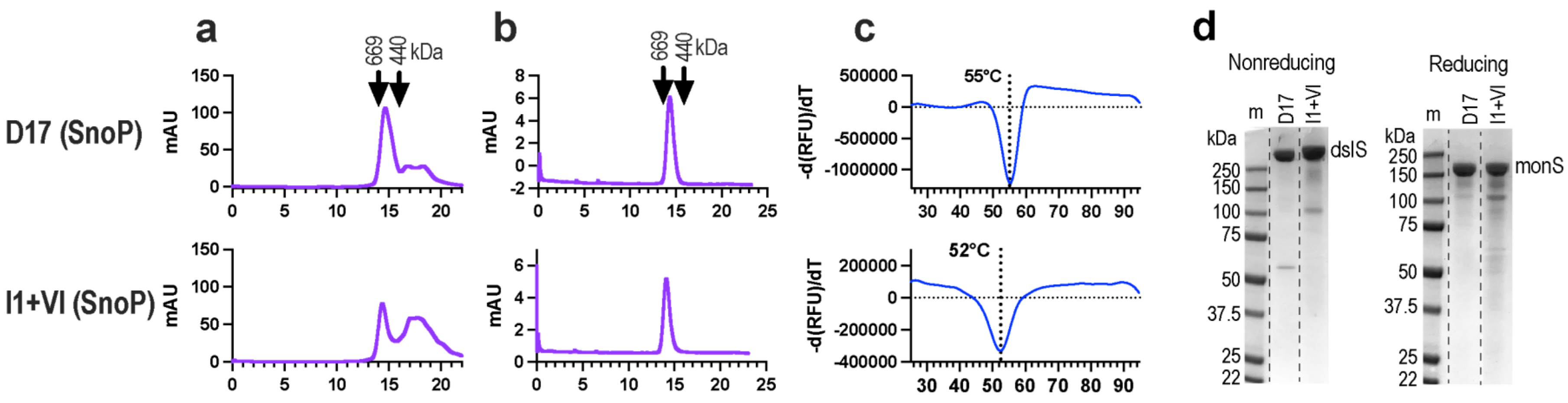


**Supplementary Fig. 4.** Effects of reverting prefusion-stabilizing proline mutations on the biochemical characteristics of spike oligomers truncated at N1192. **a.** Superose 6 SEC of secreted spike proteins eluted from TALON resin. **b.** Superose 6 SEC of purified oligomers from A following a freeze (-80°C)-thaw cycle. **c.** Melting temperature of purified spike oligomers shown in **b** determined by DSF. **d.** SDS-PAGE of purified spike proteins shown in **b**. The lanes were reorganized from single nonreducing and reducing gels (dashed lines).

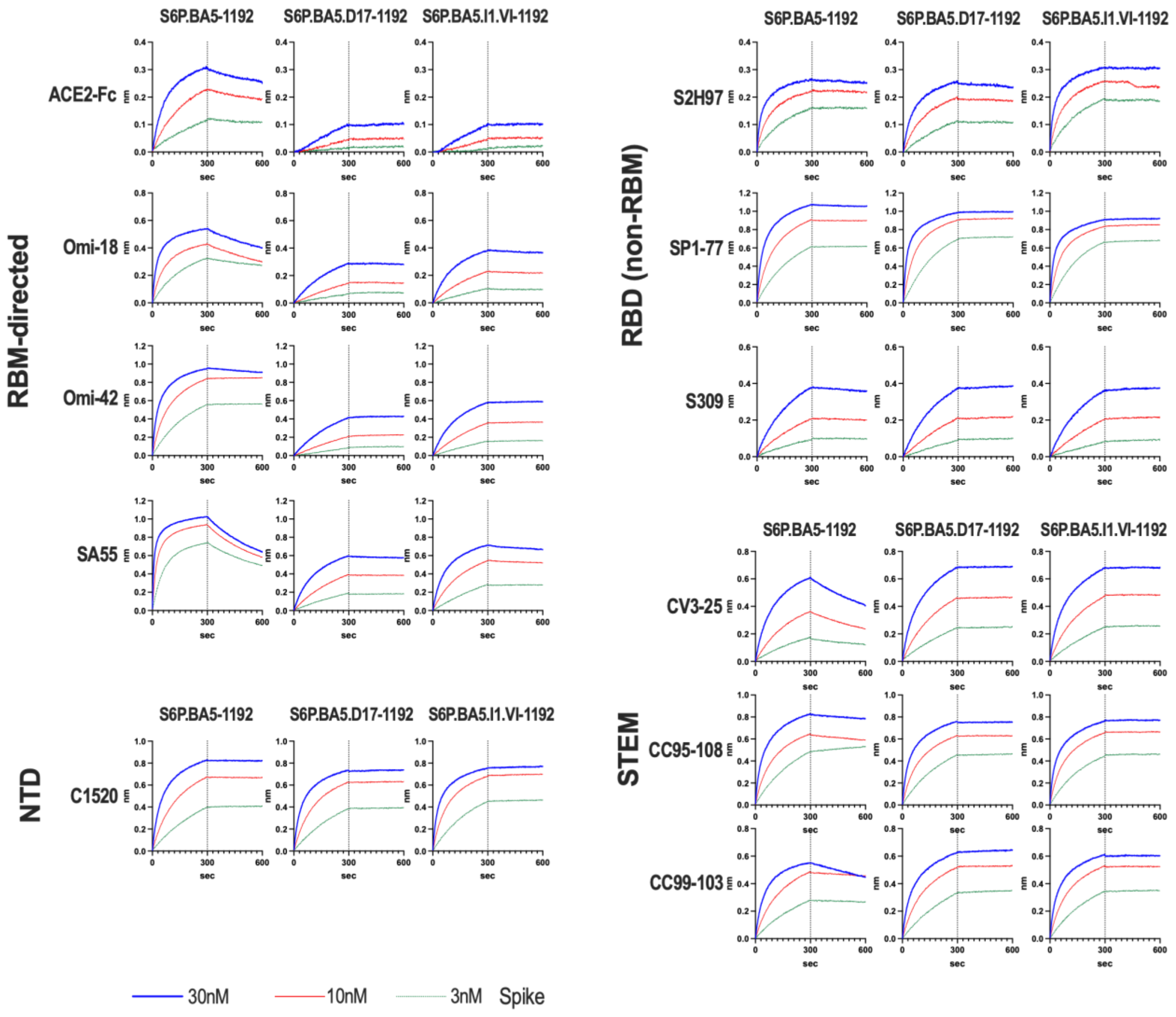


**Supplementary Fig. 5.** Epitope profiles of purified omicron BA.5 spike oligomers with D17 and I1 mutations determined in BLI with neutralizing ligands immobilized on anti-human IgG Fc capture biosensors. Association phase: 0-300 sec; dissociation phase: 301-600 sec.

**Supplementary Table 2. Binding kinetics of S2P.BA5-1192 and S6P.BA5-1192 spike constructs to ACE2-Fc and human monoclonal NAbs**

| Ligand | Spike analyte | Req | KD (M) | Ka (1/Ms) | Kdis (1/s) | $R^2$ |
|---|---|---|---|---|---|---|
| C1520 | S6P.BA5-1192 | 0.6693 | <1.0x10$^{-12}$ | 5.7x10$^{-5}$ | 4.1x10$^{-7}$ | 0.9932 |
| | +D17 | 0.6246 | <1. 0x10$^{-12}$ | 6.9 x10$^{-5}$ | 4.7 x10$^{-7}$ | 0.9914 |
| | +I1.VI | 0.677 | <1. 0x10$^{-12}$ | 9.0 x10$^{-5}$ | 3.7 x10$^{-7}$ | 0.9831 |
| ACE2-Fc | S6P.BA5-1192 | 0.2261 | 1.3x10$^{-9}$ | 4.5 x10$^{-5}$ | 5.8 x10$^{-4}$ | 0.9974 |
| | +D17 | 0.0435 | 7.6x10$^{-12}$ | 3.6 x10$^{-4}$ | 2.7 x10$^{-7}$ | 0.9911 |
| | +I1.VI | 0.0404 | 1.2x10$^{-11}$ | 3.5 x10$^{-4}$ | 4.2 x10$^{-7}$ | 0.9835 |
| Omi-18 | S6P.BA5-1192 | 0.4243 | 8.5x10$^{-10}$ | 1.1 x10$^{-6}$ | 9.2 x10$^{-4}$ | 0.9841 |
| | +D17 | 0.1436 | 8.1 x10$^{-11}$ | 1.6 x10$^{-5}$ | 1.3 x10$^{-5}$ | 0.999 |
| | +I1.VI | 0.226 | 5.8 x10$^{-10}$ | 2.8 x10$^{-5}$ | 1.6 x10$^{-4}$ | 0.999 |
| Omi-42 | S6P.BA5-1192 | 0.8375 | 6. 0x10$^{-12}$ | 8.7 x10$^{-5}$ | 5.4 x10$^{-6}$ | 0.9886 |
| | +D17 | 0.2047 | 3.6x10$^{-12}$ | 9.6 x10$^{-4}$ | 3.4 x10$^{-7}$ | 0.9983 |
| | +I1.VI | 0.3507 | 1.5x10$^{-12}$ | 2.4 x10$^{-5}$ | 3.6 x10$^{-6}$ | 0.9993 |
| SA55 | S6P.BA5-1192 | 0.9363 | 4.7 x10$^{-10}$ | 3.1x10$^{-6}$ | 1.5 x10$^{-3}$ | 0.9709 |
| | +D17 | 0.3886 | 3.2 x10$^{-10}$ | 3.0 x10$^{-5}$ | 9.6 x10$^{-5}$ | 0.9986 |
| | +I1.VI | 0.5395 | 3.5 x10$^{-10}$ | 4.6 x10$^{-5}$ | 1.6 x10$^{-4}$ | 0.9971 |
| S2H97 | S6P.BA5-1192 | 0.2213 | 1.2 x10$^{-11}$ | 1.1x10$^{-6}$ | 1.4 x10$^{-5}$ | 0.9798 |
| | +D17 | 0.1989 | 3.4 x10$^{-10}$ | 5.6 x10$^{-5}$ | 1.9 x10$^{-4}$ | 0.9933 |
| | +I1.VI | 0.2555 | 2.9 x10$^{-11}$ | 1.2x10$^{-6}$ | 3.4 x10$^{-5}$ | 0.9689 |
| SP1-77 | S6P.BA5-1192 | 0.9057 | <1.0x10$^{-12}$ | 8.5 x10$^{-5}$ | 5.9 x10$^{-6}$ | 0.9843 |
| | +D17 | 0.9031 | <1.0x10$^{-12}$ | 1.2 x10$^{-6}$ | 4.2 x10$^{-6}$ | 0.9797 |
| | +I1.VI | 0.8306 | <1.0x10$^{-12}$ | 1.2 x10$^{-6}$ | 4.5 x10$^{-6}$ | 0.9751 |
| S309 | S6P.BA5-1192 | 0.2034 | 9.4 x10$^{-10}$ | 2.0 x10$^{-5}$ | 1.8 x10$^{-4}$ | 0.9988 |
| | +D17 | 0.2088 | 2.5x10$^{-12}$ | 1.6 x10$^{-5}$ | 4.1 x10$^{-6}$ | 0.999 |
| | +I1.VI | 0.2007 | 1.8x10$^{-12}$ | 1.5 x10$^{-5}$ | 2.8 x10$^{-6}$ | 0.9988 |
| CV3-25 | S6P.BA5-1192 | 0.3602 | 4.4 x10$^{-9}$ | 3.1 x10$^{-5}$ | 1.4 x10$^{-3}$ | 0.9982 |
| | +D17 | 0.4558 | 1.5x10$^{-12}$ | 3.4 x10$^{-5}$ | 5.4 x10$^{-6}$ | 0.9965 |
| | +I1.VI | 0.4697 | 1.1x10$^{-12}$ | 3.8 x10$^{-5}$ | 4.1 x10$^{-6}$ | 0.9957 |
| CC99 | S6P.BA5-1192 | 0.4832 | 6.6 x10$^{-10}$ | 6.1 x10$^{-5}$ | 4.1 x10$^{-4}$ | 0.9949 |
| -103 | +D17 | 0.5137 | <1.0x10$^{-12}$ | 5.4 x10$^{-5}$ | 4.3 x10$^{-6}$ | 0.9874 |
| | +I1.VI | 0.5268 | <1.0x10$^{-12}$ | 7.1 x10$^{-5}$ | 4.6 x10$^{-6}$ | 0.9859 |
| CC95 | S6P.BA5-1192 | 0.6411 | 1.2 x10$^{-10}$ | 7.2 x10$^{-5}$ | 8.5 x10$^{-5}$ | 0.9796 |
| -108 | +D17 | 0.626 | <1.0x10$^{-12}$ | 7.4 x10$^{-5}$ | 5.2 x10$^{-6}$ | 0.983 |
| | +I1.VI | 0.6524 | <1.0x10$^{-12}$ | 8.2 x10$^{-5}$ | 5.5 x10$^{-6}$ | 0.9764 |

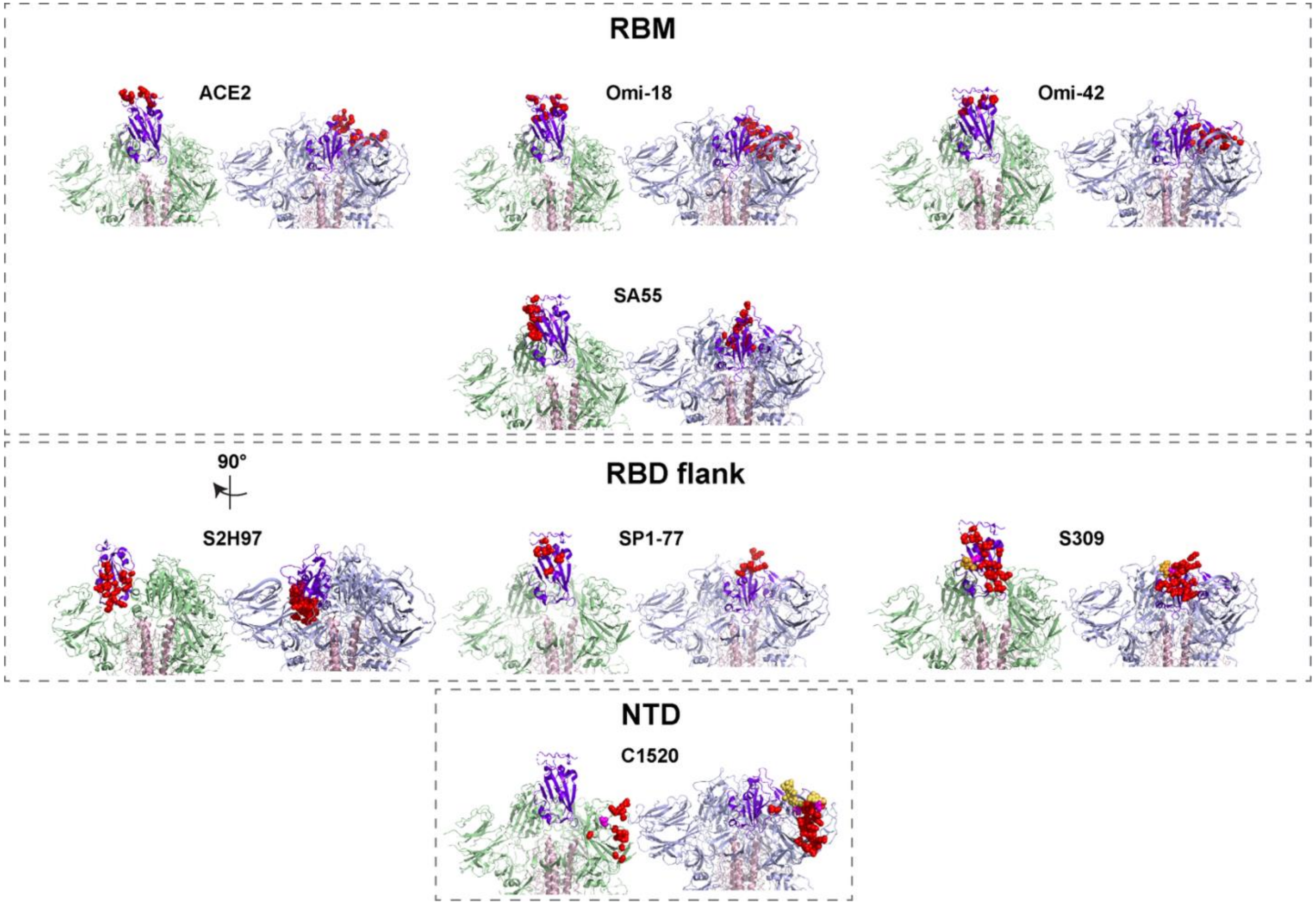


**Supplementary Fig. 6.** Location of mNAb epitopes on the SARS CoV-2 RBD when in the 1RBD-up open (left) and 3RBD-down closed (right) conformation. Amino acids (red) and glycans (orange) forming the RBM and various mNAb epitopes are shown in CPK on 1 RBD (purple). Drawn with PyMOL using the PDB IDs 6VSB (1RBD-up, open) and 6XR8 (3RBD-down, closed).

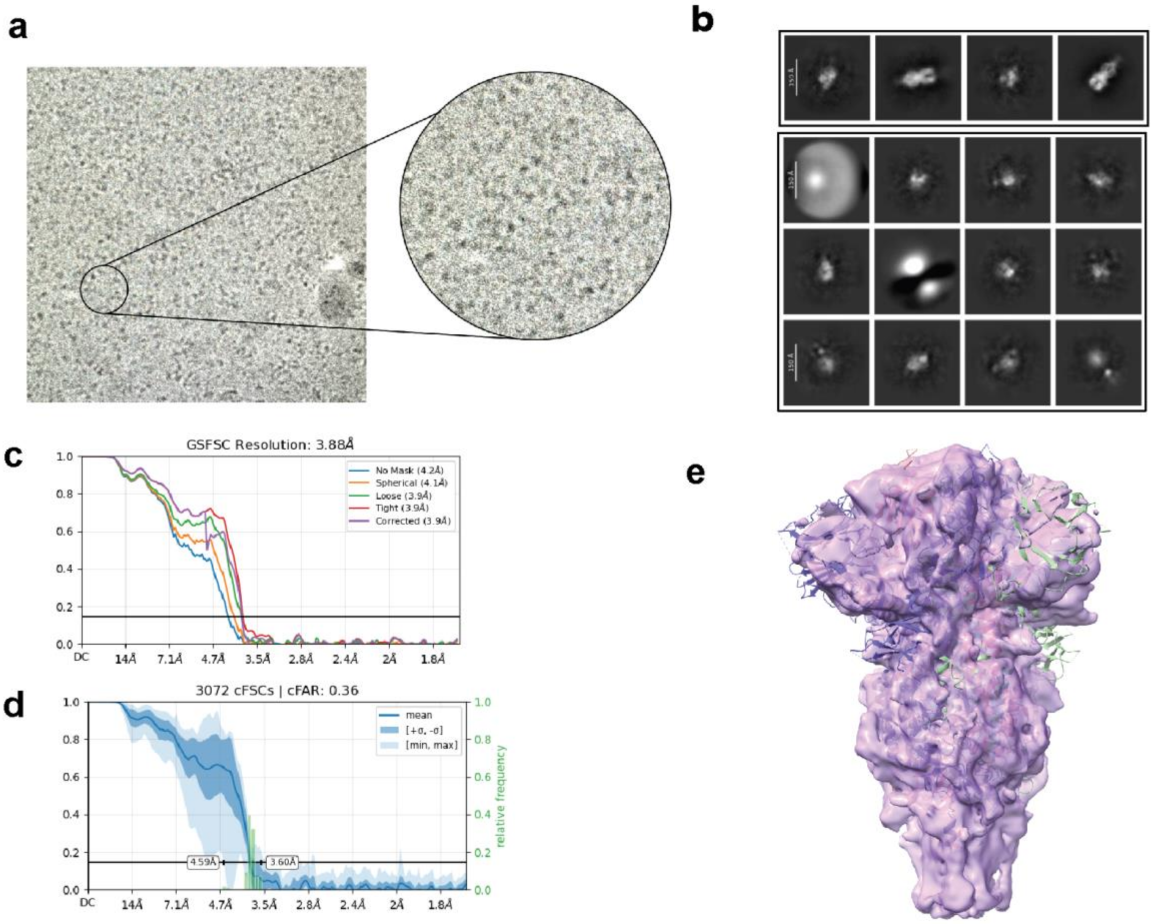


**Supplementary Fig. 7.** Cryo-EM reconstruction of S6P.BA286-1192.D17 trimer. **a.** Cryo-electronmicrograph collected from Tecnai Spirit G2 TEM (FEI); **b.** 2D class averages showing the pre-fusion conformation (row 1) and “junk” classes (rows 2-4). No evidence for the presence of post-fusion conformation was found; **c.** Fourier Shell Correlation (FSC) curves of the non-uniform refinement presented in E; **d.** The cFSC analysis shows strong anisotropy resulting from a lack of top views of the trimer; **e.** The final map obtained from non-uniform refinement, fitted with PDB structure 6X2C.

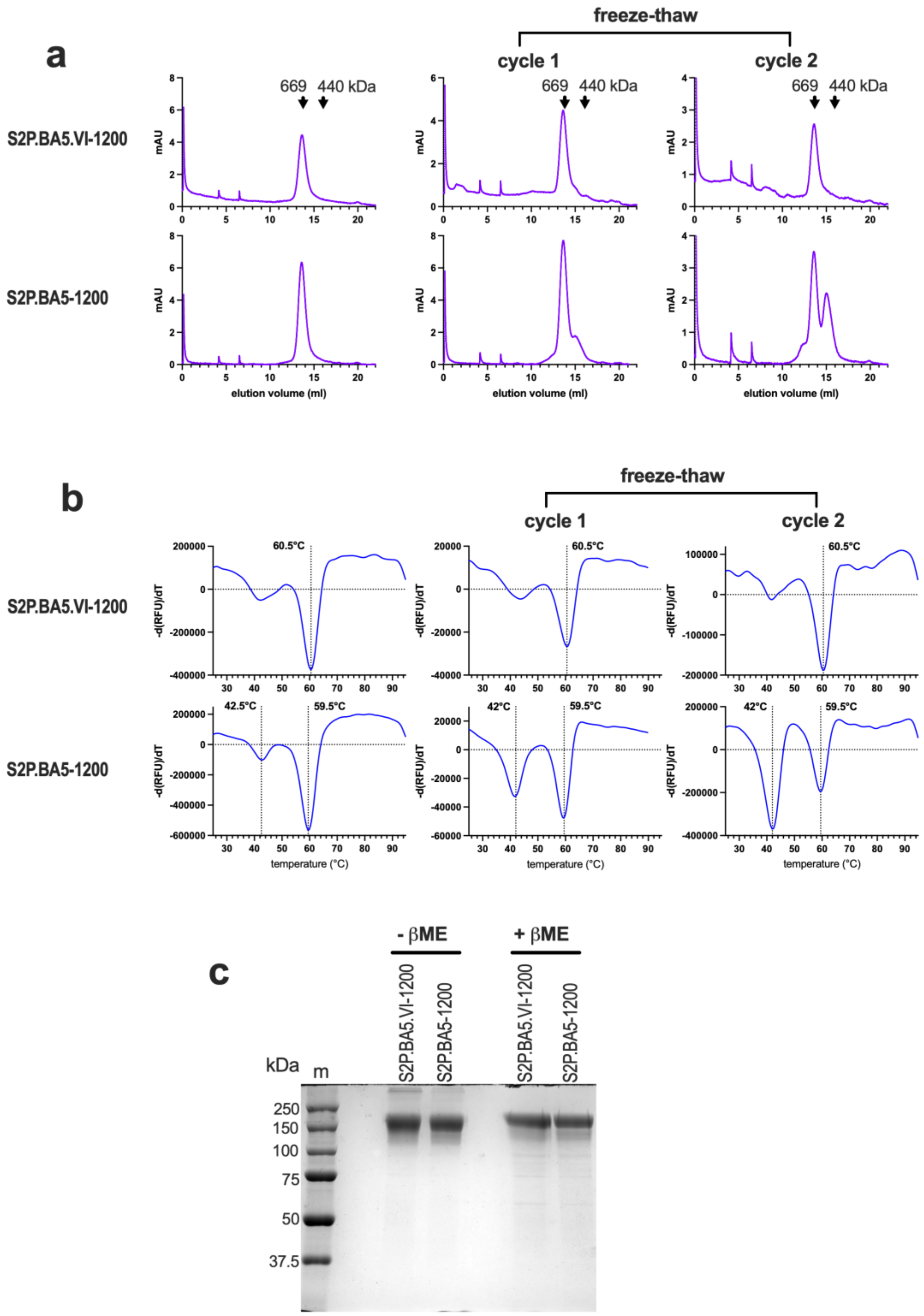


**Supplementary Fig. 8.** Biochemical characteristics of S2P.BA5 oligomers truncated at L1200 with and without the VI mutation. **a.** *Left*: SEC immediately after initial TALON and SEC purification; *middle and right:* SEC after additional 1 and 2 freeze (-80°C)-thaw cycles, respectively. **b.** *Left*: DSF immediately after initial TALON and SEC purification; *middle and right:* DSF after additional 1 and 2 freeze (-80°C)-thaw cycles, respectively. **c.** SDS-PAGE of purified spike oligomers.

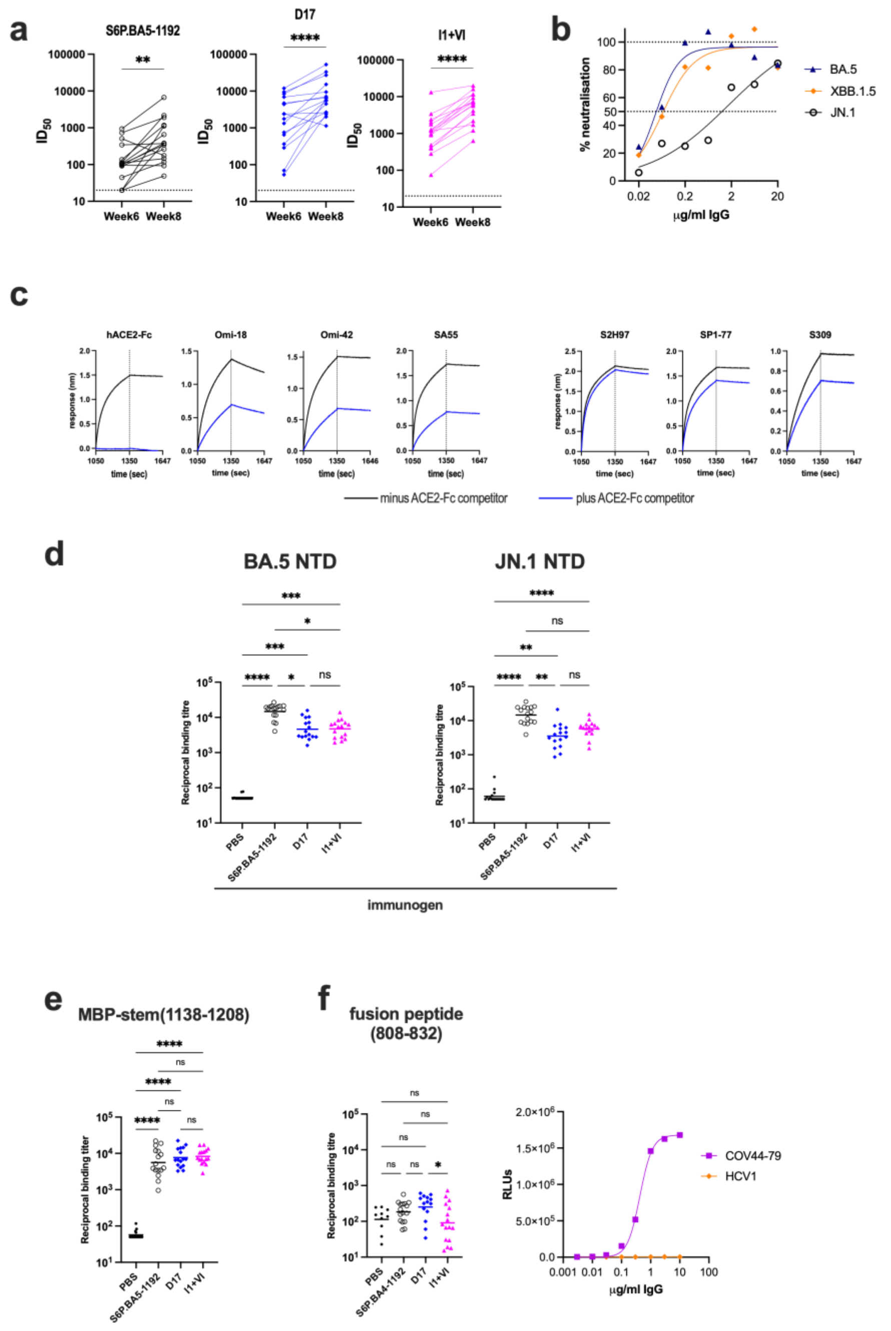


**Supplementary Fig. 9. a.** Boosting effect of 3rd dose with respect to homologous neutralizing antibody titres. **, P<0.01; ****, P<0.0001, Wilcoxon rank test. **b.** Neutralization of SARS CoV-2 variants by mNAb Omi-42. **c.** ACE2-Fc blockade of neutralizing ligands binding to biotin-omicron BA.5 RBD attached to streptavidin biosensors in BLI. The ligands are indicated above the sensograms. **d.** NTD binding titres of sera obtained after 3 immunizations with experimental vaccines. The variant NTDs are indicated above the graphs. **e.** Recombinant MBP-stem(1138-1208) binding titres of sera obtained after 3 immunizations with experimental vaccines. **f.** Fusion peptide binding titres of sera obtained after 3 immunizations with experimental vaccines. *Right panel:* Binding of anti-fusion peptide mNAb COV44-79 or HCV E2-specific mNAb HCV1 to fusion peptide. For **d**, **e**, **f**, horizontal bars are the geometric means; *, $P < 0.05$; **, $P < 0.01$: ***, $P < 0.001$; ****, $P < 0.0001$, Kruskal-Wallis test.

### Supplementary Material

**a**

| | | subdomain 1 (SD1) ← → 570 / 571 | heptad repeat 1 (HR1) ← → 967 | % AA ID BA.5 |
|---|---|---|---|---|
| **Clade 3** | BM4831 | ...TNSTKKFQPFQQFGRDV**SD**FTDSVRDPKTLEILDIAP | ...TLVKQL**S**SNFGAISSVLNDILSRLDKV... | 71.2 |
| | PRD0038 | ...TSSTKKFQPFQQFGRDV**SD**FTDSVRDPKTLEILDISP | ...TLVKQL**S**SNFGAISSVLNDILSRLDKV... | 72.5 |
| | RfGB02 | ...TDSVKKFQPFQQFGRDS**SD**FTDSVKDPKTLEILDITP | ...TLVKQL**S**SNFGAISSVLNDILSRLDKV... | 72.2 |
| | Khosta2 | ...TDSNKKFQPFQQFGRDS**AD**FTDSVKDPKTLEILDITP | ...TLVKQL**S**SNFGAISSVLNDILSRLDKV... | 71.1 |
| **Clade 1b** | ZXC21 | ...TDSSKRFQSFQQFGKDA**SD**FIDSVRDPQTLEILDITP | ...TLVKQL**S**SNFGAISSVLNDILSRLDKV... | 79.9 |
| | PangoMP789 | ...TESSKKFLPFQQFGRDI**AD**TTDAVRDPQTLEILDITP | ...TLVKQL**S**SNFGAISSVLNDILSRLDKV... | 88.9 |
| | PangoP1E | ...TTSKKQFLPFQQFGRDI**SD**TTDAVRDPQTLEILDITP | ...TLVKQL**S**SNFGAISSVLNDILSRLDKV... | 91.0 |
| | BA286 | ...TKSNKKFLPFQQFGRDI**VD**TTDAVRDPQTLEILDITP | ...TLVKQL**S**SKFGAISSVLNDILSRLDKV... | 97.3 |
| | BA5 | ...TESNKKFLPFQQFGRDI**AD**TTDAVRDPQTLEILDITP | ...TLVKQL**S**SKFGAISSVLNDILSRLDKV... | 100 |
| **Clade 1a** | LYRa3 | ...TPSSKRFQPFQQFGRDV**SD**FTDSVRDPKTLEVLDISP | ...TLVKQL**S**SNFGAISSVLNDILSRLDKV... | 75.7 |
| | BtSY1 | ...TPSSKRFQPFQQFGRDV**SD**FTDSVRDPKTSEILDISP | ...TLVKQL**S**SNFGAISSVLNDILSRLDKV... | 75.6 |
| | WIV1 | ...TPSSKRFQPFQQFGRDV**SD**FTDSVRDPKTSEILDISP | ...TLVKQL**S**SNFGAISSVLNDILSRLDKV... | 76.2 |
| | SARSGD01 | ...TPSSKRFQPFQQFGRDV**SD**FTDSVRDPKTSEILDISP | ...TLVKQL**S**SNFGAISSVLNDILSRLDKV... | 75.5 |
| **Clade 2** | HKU3 | ...TSSSKRFQSFQQFGRDT**SD**FTDSVRDPQTLEILDISP | ...TLVKQL**S**SNFGAISSVLNDILSRLDKV... | 76.5 |
| | RpShaanxi11 | ...TASSKKFQSFQQFGRDA**SD**FTDSVRDPQTLEILDISP | ...TLVKQL**S**SNFGAISSVLNDILSRLDKV... | 75.3 |
| | CpYunnan11 | ...TDSSKKFQSFQQFGRDA**SD**FTDSVRDPQTLQILDISP | ...TLVKQL**S**SNFGAISSVLNDILSRLDKV... | 75.1 |
| | | * * *:* *****:* * *:*:**:* ::***:* | ********:****************** | |

**Supplementary Fig. 10. a.** Alignment of Sarbecovirus spike SD1 and HR1 fragment sequences to identify A570, D571 and S967-homologous amino acids for Cys substitution mutagenesis.

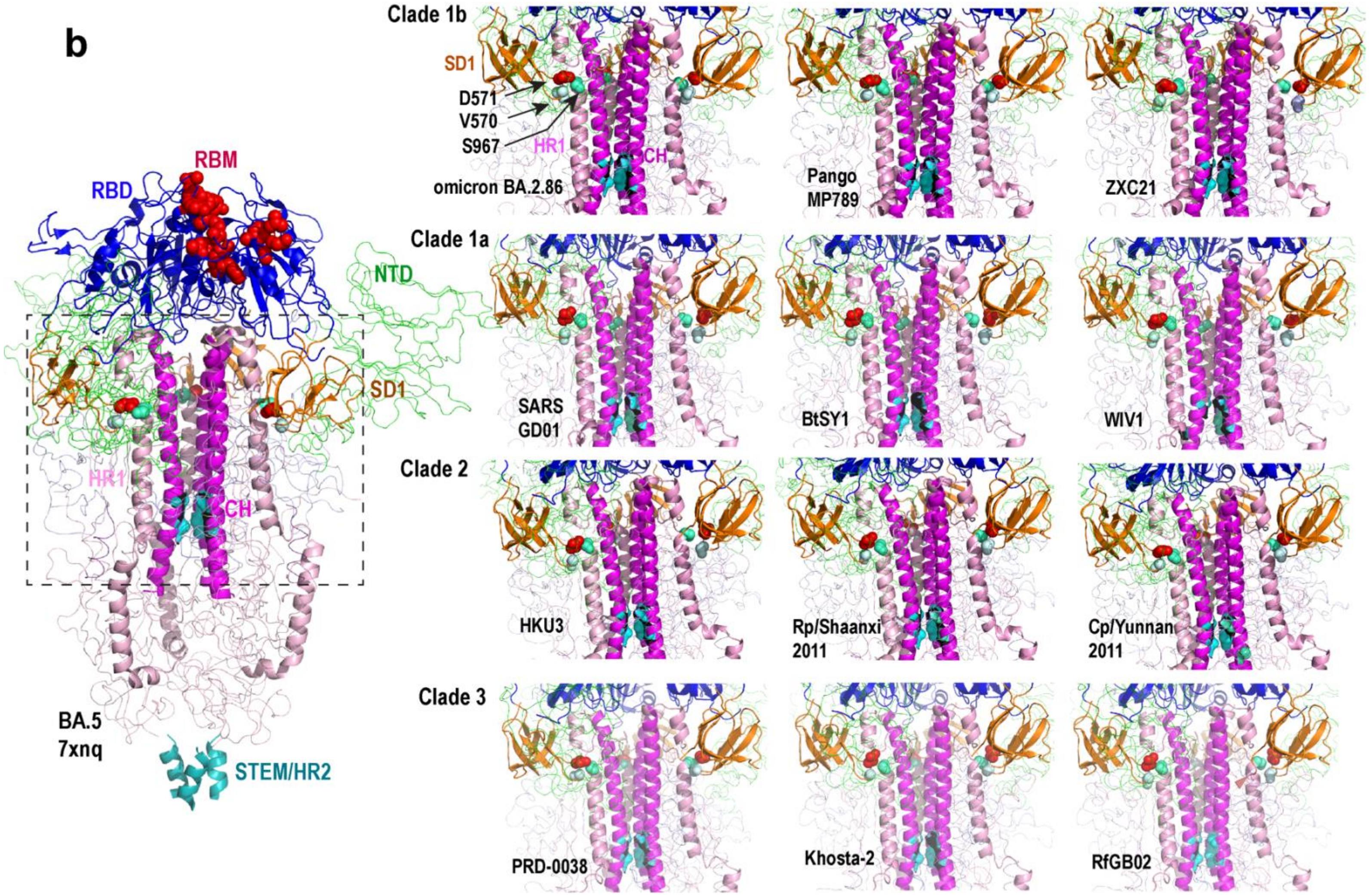


**Supplementary Fig. 10. b.** AlphaFold 3-derived homology models of Sarbecovirus spike trimers, focussing on the region around the A570, D571 and S967 triad. Sarbecovirus spike sequences corresponding to SARS CoV-2 Hu-1 amino acids 16-1192 were inputted to the AlphaFold3 portal. Model 0 is shown in all cases.

**a**

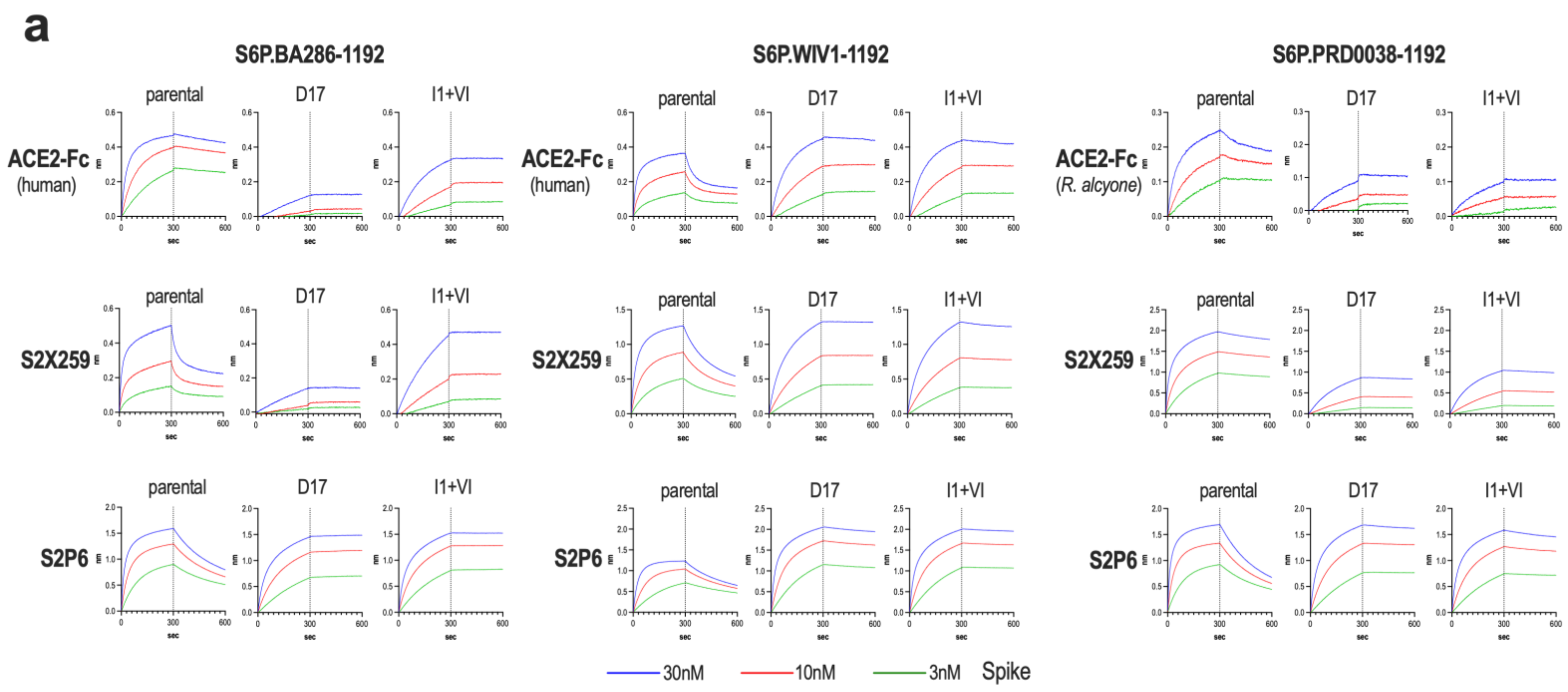


**b**

| | | S6P.BA286-1192 | | | | S6P.WIV1-1192 | | | | S6P.PRD-1192 | | | |
|---|---|---|---|---|---|---|---|---|---|---|---|---|---|
| | | R300/30nM[a] (nm) | KD (M) | ka (1/Ms) | kd (1/s) | R300/30nM (nm) | KD (M) | ka (1/Ms) | kd (1/s) | R300/30nM (nm) | KD (M) | ka (1/Ms) | kd (1/s) |
| ACE2-Fc | parental | 0.47 | 2.7E-10 | 9.2E05 | 2.4E-04 | 0.36 | 2.2E-09 | 1.5E06 | 3.2E-03 | 0.2 | 1.2E-09 | 5.5E05 | 6.3E-04 |
| | D17 | 0.12 | 9.6E-12 | 3.5E04 | 3.4E-07 | 0.45 | 1.3E-12 | 3.4E05 | 4.4E-07 | 0.1 | 7.1E-12 | 1.4E05 | 2.6E-07 |
| | I1.VI | 0.32 | 2.4E-12 | 1.4E05 | 3.5E-07 | 0.40 | 5.9E-11 | 3.7E05 | 2.2E-05 | 0.1 | 2.9E-12 | 3.7E04 | 4.1E-07 |
| S2X259 | parental | 0.50 | 2.0E-09 | 1.5E06 | 3.1E-03 | 1.3 | 3.2E-09 | 9.6E05 | 3.1E-03 | 2.0 | 2.4E-10 | 9.6E05 | 2.3E-04 |
| | D17 | 0.14 | 7.2E-12 | 3.6E04 | 2.6E-07 | 1.3 | <1.0E-12 | 3.2E05 | 2.9E-07 | 0.9 | 4.8E-10 | 1.7E05 | 8.0E-05 |
| | I1.VI | 0.45 | 5.0E-12 | 6.4E04 | 3.2E-07 | 1.3 | 4.1E-10 | 3.2E05 | 1.3E-04 | 1.0 | 5.1E-10 | 2.4E05 | 1.E-04 |
| S2P6 | parental | 1.6 | 1.6E-09 | 1.5E06 | 2.3E-03 | 1.2 | 1.8E-09 | 1.1E06 | 2.1E-03 | 1.7 | 1.7E-09 | 1.7E06 | 3.0E-03 |
| | D17 | 1.5 | <1.0E-12 | 5.0E05 | 4.5E-07 | 2.0 | 1.3E-10 | 9.2E05 | 1.1E-04 | 1.7 | 7.3E-11 | 5.9E05 | 4.3E-05 |
| | I1.VI | 1.5 | <1.0E-12 | 6.1E05 | 5.0E-07 | 2.0 | 6.5E-12 | 7.4E05 | 4.8E-06 | 1.6 | 2.9E-10 | 6.8E05 | 2.0E-04 |

[a]Response at 300 sec, 30 nM spike analyte

**Supplementary Fig. 11. a.** Binding of Sarbecovirus S6P-1192 spike oligomers to ACE2 and neutralizing human mNAbs in BLI. **b.** Binding kinetics from **a**.

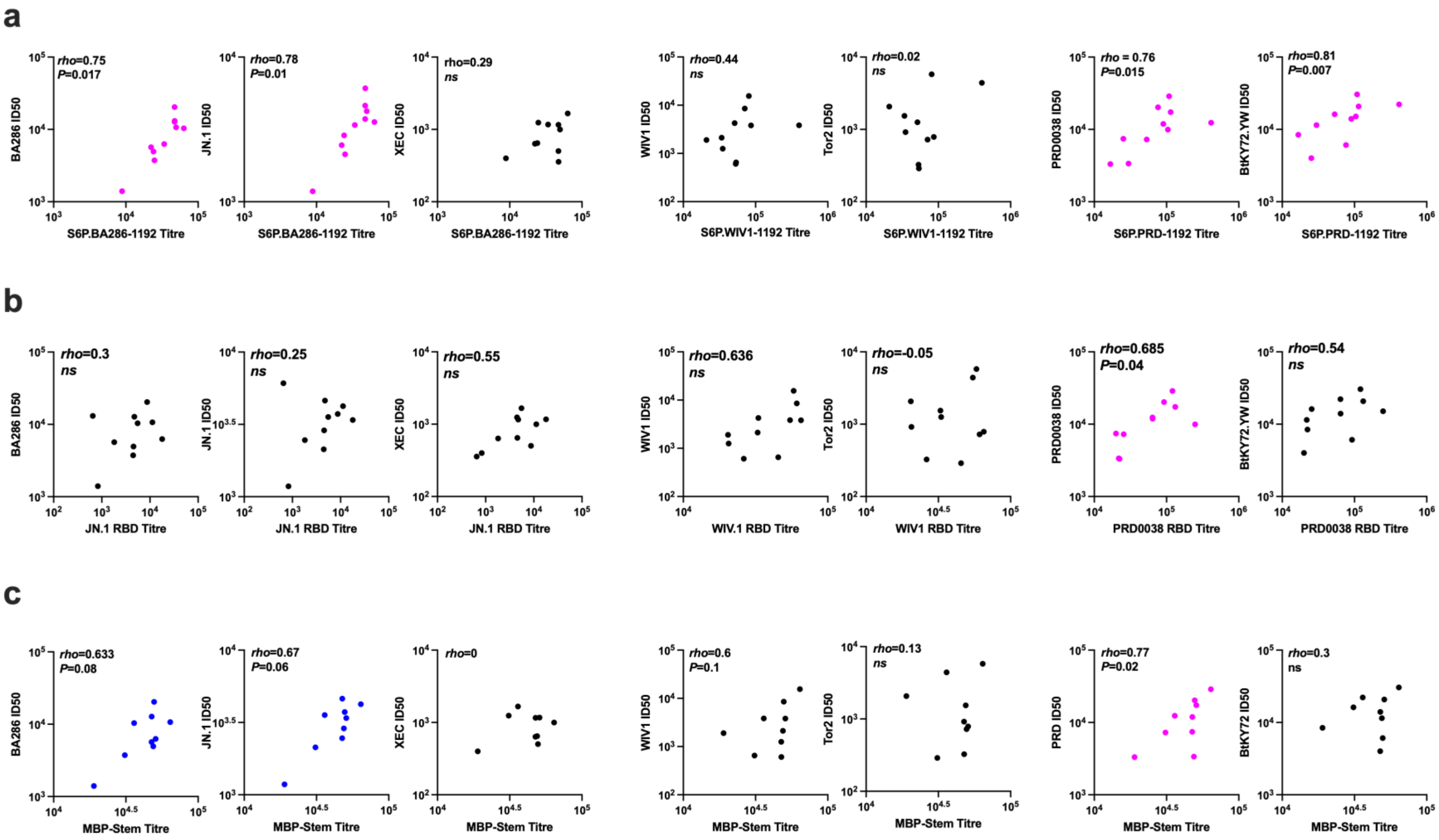

**Supplementary Fig. 12**. Correlations between neutralization ID50 and, **a,** S6P-1192 binding titre, **b,** RBD binding titre, **c,** MBP-stem(1138-1208) binding titre in D17-Trivalent immune sera. Data showing significant correlations are highlighted in magenta. Spearman *rho* and *P* values were determined in GraphPad Prism v9.3.0. Blue points: *rho* suggests moderate correlation but *P*>0.05.

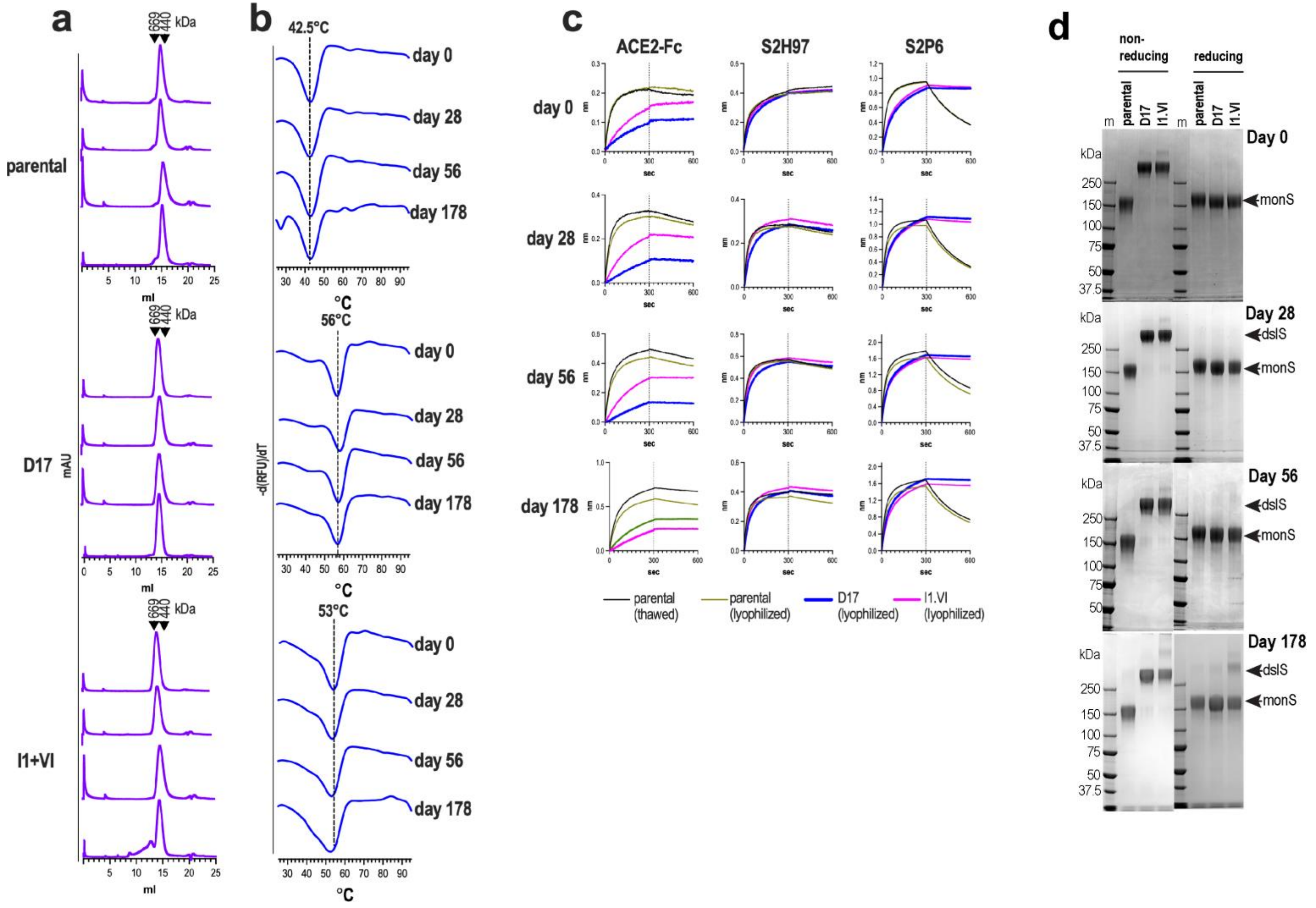


**Supplementary Fig. 13.** Biochemical and antigenic analysis of purified parental, D17 and I1+VI -mutated S6P.BA286-1192 oligomers following lyophilization and storage at ambient temperature for up to 178 days. **a.** Superose 6 SEC. **b.** DSF. **c.** Reactivity with ACE2-Fc and mNAbs in BLI. **d.** SDS-PAGE under non-reducing and reducing conditions. The gels were stained with Coomassie brilliant blue dye. dslS, disulfide linked S; monS, S monomers

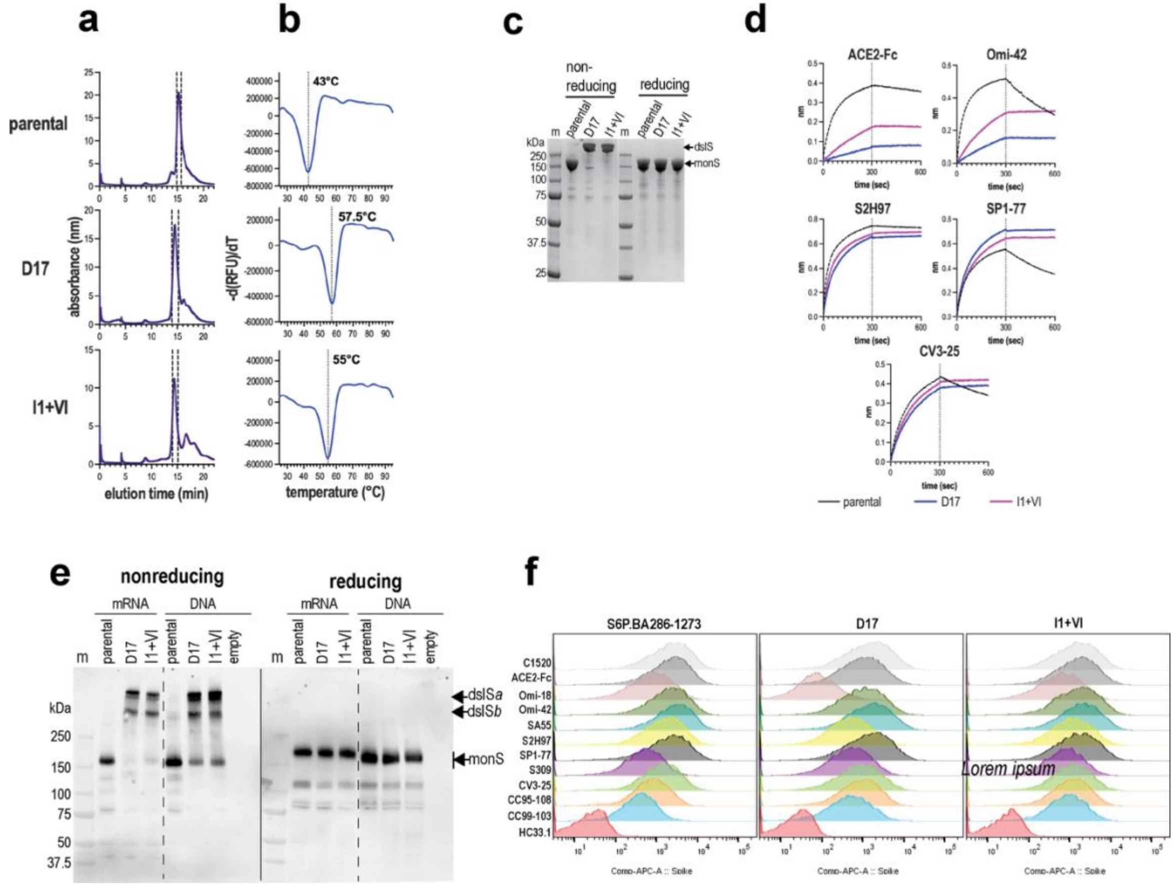


**Supplementary Fig. 14.** Biochemical and antigenic characteristics of S6P.BA286-1192 and S6P.BA286-1273 glycoproteins expressed from mRNA. **a.** Superose 6 SEC of secreted S6P.BA286-1192 spike variants eluted from TALON resin. Fractions within the verticle dashed lines were pooled and analysed in B, C and D. **b.** Melting temperatures of purified S6P.BA286-1192 trimers determined by DSF. **c.** SDS-PAGE of purified Spike oligomers. dslS, disulfide-linked spike; monS; spike monomer. **d.** Epitope profiles of purified omicron BA.5 spike oligomers with D17 and I1 mutations determined in BLI with neutralizing ligands immobilized on anti-human IgG Fc capture biosensors. **e.** Expression of full-length S6P.BA286-1273 spikes from mRNA or pcDNA3 expression vectors detected by SDS-PAGE/Western blotting of transfected 293T cell lysates. dslS*a*, dslS*b*, disulfide-bonded spike; monS, monomeric spike. **f.** Flow cytometry of mRNA-expressed S6P.BA286-1273 spikes on the surface of transfected 293T cells using neutralizing ligands.
